# Multi-omics integration and regulatory inference for unpaired single-cell data with a graph-linked unified embedding framework

**DOI:** 10.1101/2021.08.22.457275

**Authors:** Zhi-Jie Cao, Ge Gao

## Abstract

With the ever-increasing amount of single-cell multi-omics data accumulated during the past years, effective and efficient computational integration is becoming a serious challenge. One major obstacle of unpaired multi-omics integration is the feature discrepancies among omics layers. Here, we propose a computational framework called GLUE (graph-linked unified embedding), which utilizes accessible prior knowledge about regulatory interactions to bridge the gaps between feature spaces. Systematic benchmarks demonstrated that GLUE is accurate, robust and scalable. We further employed GLUE for various challenging tasks, including triple-omics integration, model-based regulatory inference and multi-omics human cell atlas construction (over millions of cells) and found that GLUE achieved superior performance for each task. As a generalizable framework, GLUE features a modular design that can be flexibly extended and enhanced for new analysis tasks. The full package is available online at https://github.com/gao-lab/GLUE for the community.

## Introduction

Recent technological advances in single-cell sequencing have enabled the probing of regulatory maps through multiple omics layers, such as chromatin accessibility (scATAC-seq^1, 2^), DNA methylation (snmC-seq^3^, sci-MET^4^) and the transcriptome (scRNA-seq^5, 6^), offering a unique opportunity to unveil the underlying regulatory bases for the functionalities of diverse cell types^7^. While simultaneous assays are emerging recently^8-11^, different omics are usually measured independently and produce unpaired data, which calls for effective and efficient *in silico* multi-omics integration^12, 13^.

Computationally, one major obstacle faced when integrating unpaired multi-omics data is the distinct feature spaces of different modalities (e.g., accessible chromatin regions in scATAC-seq vs. genes in scRNA-seq)^14^. A quick fix is to convert multimodality data into one common feature space based on prior information and apply single-omics data integration methods^15-17^. Such explicit “feature conversion” is straightforward, but has been reported to result in significant information loss^18^. Algorithms based on coupled matrix factorization circumvent explicit conversion but hardly handle more than two omics layers^19, 20^. An alternative option is to match cells from different omics layers via nonlinear manifold alignment, which removes the requirement of prior knowledge completely and could reduce inter-modality information loss in theory^21, 22^; however, this technique has mostly been applied to continuous, trajectory-like manifolds rather than atlases.

The ever-increasing volume of data is another serious challenge^23^. Recently developed technologies can routinely generate datasets at the scale of millions of cells^24-26^, whereas current integration methods have only been applied to datasets with much smaller volumes^15, 17, 19-22^. To catch up with the growth in data throughput, computational integration methods should be designed with scalability in mind.

Hereby, we introduce GLUE (graph-linked unified embedding), a modular framework for integrating unpaired single-cell multi-omics data and inferring regulatory interactions simultaneously. By modeling the regulatory interactions across omics layers explicitly, GLUE bridges the gaps between various omics-specific feature spaces in a biologically intuitive manner. Systematic benchmarks and case studies demonstrate that GLUE is accurate, robust and scalable for heterogeneous single-cell multi-omics data. Furthermore, GLUE is designed as a generalizable framework that allows for easy extension and quick adoption to particular scenarios in a modular manner. GLUE is publicly accessible at https://github.com/gao-lab/GLUE.

## Results

### Integrating unpaired single-cell multi-omics data via graph-guided embeddings

Inspired by previous works, we model cell states as low-dimensional cell embeddings learned through variational autoencoders^27, 28^. Given their intrinsic differences in biological nature and assay technology, each omics layer is equipped with a separate autoencoder that uses a probabilistic generative model tailored to the layer-specific feature space (Fig. 1, Methods).

**Fig. 1.**
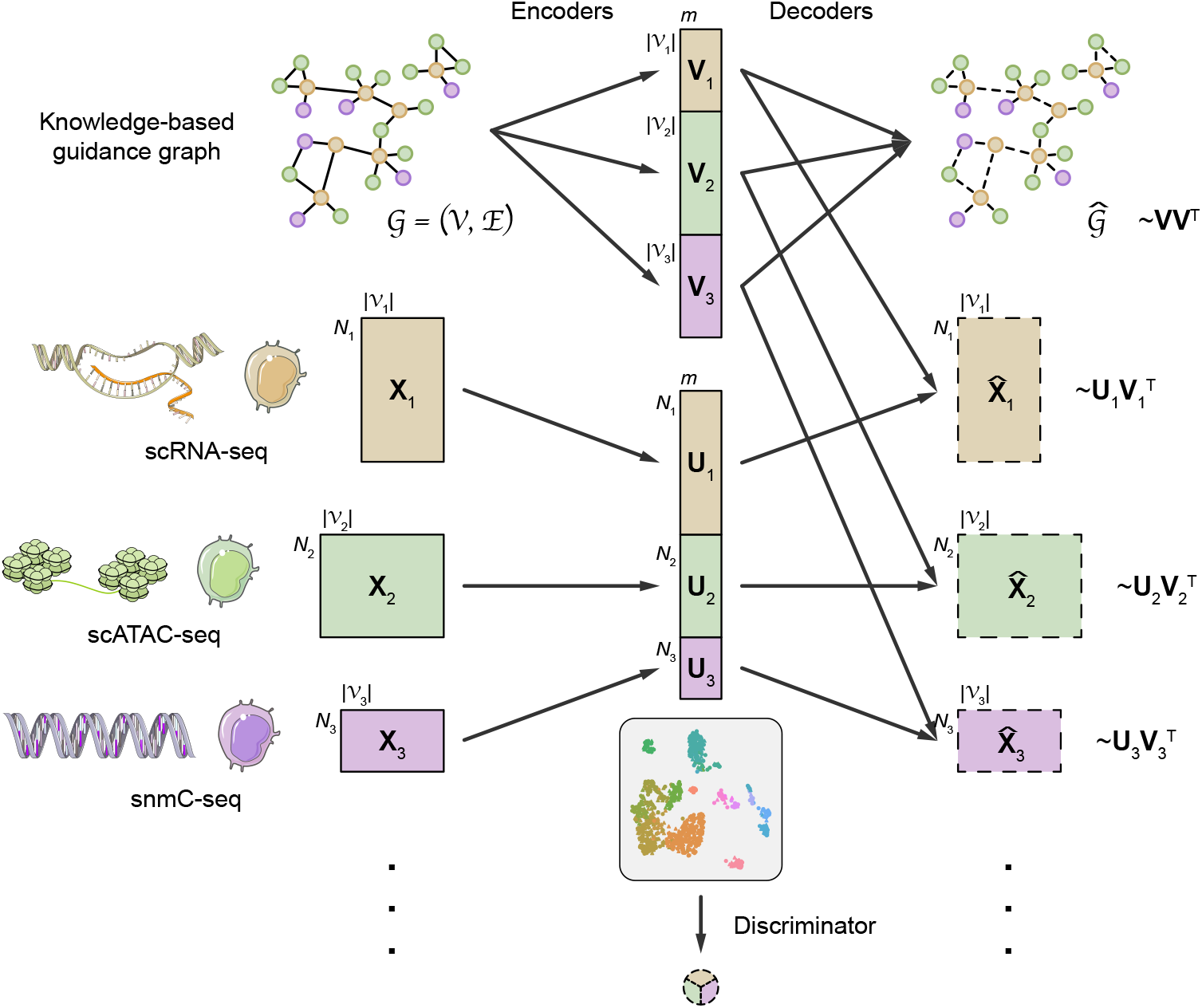
Architecture of the GLUE framework. GLUE employs omics-specific variational autoencoders to learn low-dimensional cell embeddings from each omics layer. The data dimensionality and generative distribution can differ across omics layers, but the cell embedding dimensions are shared. A graph variational autoencoder is used to learn feature embeddings from the prior knowledge-based guidance graph; these embeddings are then used as data decoder parameters. The feature embeddings effectively link the omics-specific autoencoders to ensure a consistent embedding orientation. Last, an omics discriminator is employed to align the cell embeddings of different omics layers via adversarial learning.

Taking advantage of prior biological knowledge, we propose the use of a knowledge-based graph (“guidance graph”) that explicitly models cross-layer regulatory interactions for linking layer-specific feature spaces; the vertices in the graph correspond to the features of different omics layers, and edges represent signed regulatory interactions. For example, when integrating scRNA-seq and scATAC-seq data, the vertices are genes and accessible chromatin regions (i.e., ATAC peaks), and a positive edge can be connected between an accessible region and its putative downstream gene. Then, adversarial multimodal alignment is performed as an iterative optimization procedure, guided by feature embeddings encoded from the graph^29^ (Fig. 1, Methods). Notably, when the iterative process converges, the graph can be refined with inputs from the alignment procedure and used for data-oriented regulatory inference (see below for more details).

### Systematic benchmarks demonstrate superior alignment accuracy and robustness over existing methods

We first benchmarked GLUE against multiple popular unpaired multi-omics integration methods^15-17, 21, 30^ using gold-standard datasets generated by recent simultaneous scRNA-seq and scATAC-seq technologies^8, 9, 31^.

At the cell type level, an integration method should match the corresponding cell types from different omics layers, producing cell embeddings where the cell types are clearly distinguishable and the omics layers are well mixed. Compared to other methods, GLUE achieved overall the highest cell type resolution (as quantified by mean average precision) and layer mixing (as quantified by the Seurat alignment score^32^) simultaneously (Fig. 2a); these results were also validated by UMAP visualization of the aligned cell embeddings (Supplementary Fig. 1-3).

**Fig. 2.**
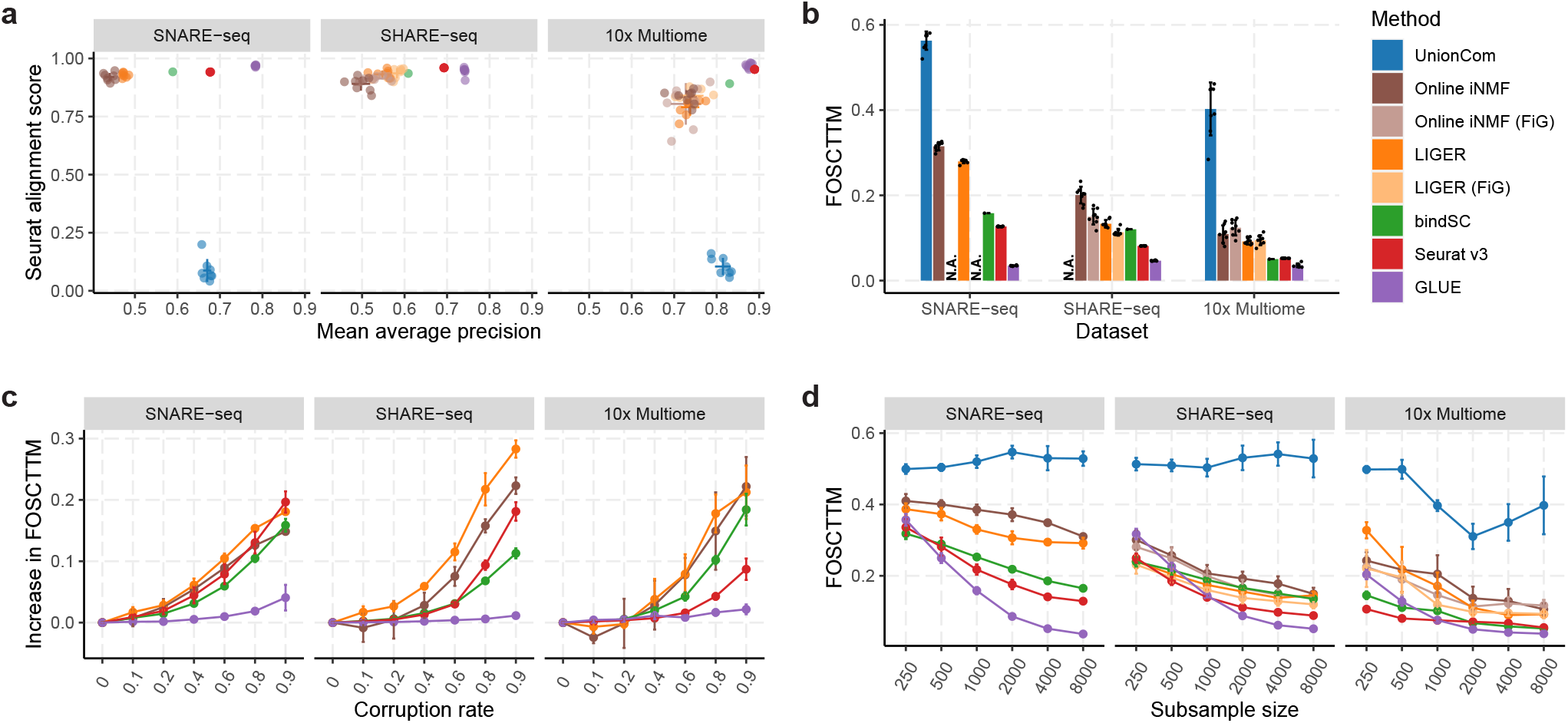
Systematic benchmarks on gold-standard datasets. **a**, Cell type resolution (quantified by mean average precision) vs. omics layer mixing (quantified by Seurat alignment score) for different integration methods. FiG (fragments in genes) is an alternative feature conversion method recommended by online iNMF and LIGER (Methods). Online iNMF and LIGER could not run with FiG conversion on the SNARE-seq data because the raw ATAC fragment file was not available. UnionCom failed to run on the SHARE-seq dataset due to memory overflow. **b**, Single-cell level alignment error (quantified by FOSCTTM) of different integration methods. **c**, Increases in FOSCTTM at different prior knowledge corruption rates for integration methods that rely on prior feature interactions. **d**, FOSCTTM values of different integration methods on subsampled datasets. The error bars indicate mean ± s.d.

An optimal integration method should produce accurate alignments not only at the cell type level but also at finer scales. Exploiting the ground truth cell-to-cell correspondence between scRNA-seq and scATAC-seq, we further quantified single-cell level alignment error via the FOSCTTM (fraction of samples closer than the true match) metric^33^. On all three datasets, GLUE achieved the lowest FOSCTTM, decreasing the alignment error by large margins compared to the second-best method on each dataset (Fig. 2b, the decreases were 3.6-fold for SNARE-seq, 1.7-fold for SHARE-seq, and 1.4-fold for 10x Multiome).

During the evaluation described above, we adopted a standard schema (ATAC peaks were linked to RNA genes if they overlapped in the gene body or proximal promoter regions) to construct the guidance graph for GLUE and to perform feature conversion for other conversion-based methods. Given that our current knowledge about the regulatory interactions is still far from prefect, a useful integration method must be robust to such inaccuracies. Thus, we further assessed the methods’ robustness to corruption of regulatory interactions by randomly replacing varying fractions of existing interactions with nonexistent ones. For all three datasets, GLUE exhibited the smallest performance changes even at high corruption rates (Fig. 2c), suggesting its superior robustness.

Given its neural network-based nature, GLUE may suffer from undertraining when working with small datasets. Thus, we repeated the evaluations using subsampled datasets of various sizes. GLUE remained the top-ranking method with as few as 2,000 cells, but the alignment error increased more steeply when the data volume decreased to less than 1,000 cells (Fig. 2d). Additionally, we also noted that the performance of GLUE was robust for a wide range of hyperparameter settings (Supplementary Fig. 4).

### GLUE enables effective triple-omics integration

Benefitting from a modular design and scalable adversarial alignment, GLUE readily extends to more than two omics layers. As a case study, we used GLUE to integrate three distinct omics layers of neuronal cells in the adult mouse cortex, including gene expression^34^, chromatin accessibility^35^, and DNA methylation^3^.

Unlike chromatin accessibility, gene body DNA methylation generally shows a negative correlation with gene expression in neuronal cells^36^. GLUE natively supports the mixture of regulatory effects by modeling edge signs in the guidance graph. Such a strategy avoids data inversion, which is required by previous methods^16, 17^ and can break data sparsity and the underlying distribution. For the triple-omics guidance graph, we linked gene body mCH and mCG levels to genes via negative edges, while the positive edges between accessible regions and genes remained the same.

The GLUE alignment successfully revealed a shared manifold of cell states across the three omics layers (Fig. 3a-d). We observed highly significant marker overlap (Fig. 3e, three-way Fisher’s exact test^37^, FDR < 2×10^−15^) for 12 out of the 14 mapped cell types (Supplementary Fig. 5, 6, Methods), indicating reliable alignment. Interestingly, we found that GLUE alignment helped improve the effects of cell typing in all omics layers, including the further partitioning of the scRNA-seq “MGE” cluster into *Pvalb*^+^ (“mPv”) and *Sst*^+^ (“mSst”) subtypes (highlighted with green circles/flows in Fig. 3, Supplementary Fig. 5), the partitioning of the scRNA-seq “CGE” cluster and scATAC-seq “Vip” cluster into *Vip*^+^ (“mVip”) and *Ndnf*^+^ (“mNdnf”) subtypes (highlighted with dark blue circles/flows in Fig. 3, Supplementary Fig. 5), and the identification of snmC-seq “mDL-3” cells and a subset of scATAC-seq “L6 IT” cells as claustrum cells (highlighted with light blue circles/flows in Fig. 3, Supplementary Fig. 5).

**Fig. 3.**
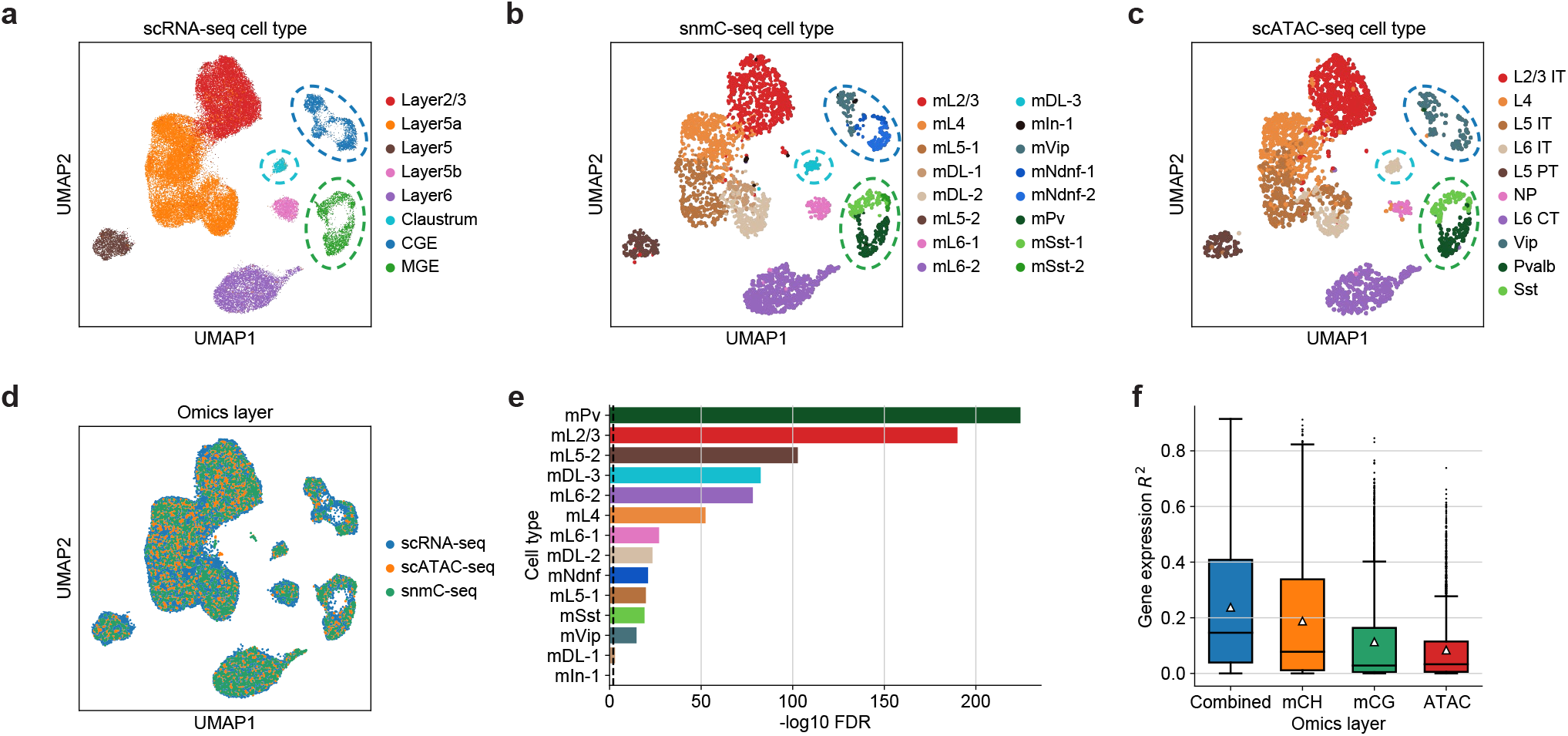
Triple-omics integration of the mouse cortex. **a**-**c**, UMAP visualizations of the integrated cell embeddings for **a**, scRNA-seq, **b**, snmC-seq, and **c**, scATAC-seq, colored by the original cell types. Cells aligning with “mPv” and “mSst” are highlighted with green circles. Cells aligning with “mNdnf” and “mVip” are highlighted with dark blue circles. Cells aligning with “mDL-3” are highlighted with light blue circles. **d**, UMAP visualizations of the integrated cell embeddings for all cells, colored by omics layers. **e**, Significance of marker gene overlap for each cell type across all three omics layers (three-way Fisher’s exact test^37^). The dashed vertical line indicates that FDR = 0.01. We observed highly significant marker overlap (FDR < 2×10^−15^) for 12 out of the 14 cell types, indicating reliable alignment. For the remaining 2 cell types, “mDL-1” had marginally significant marker overlap with FDR = 0.001, while the “mIn-1” cells in snmC-seq did not properly align with the scRNA-seq or scATAC-seq cells. **f**, Coefficient of determination (*R*^2^) for predicting gene expression based on each epigenetic layer as well as the combination of all layers. The box plots indicate the medians (centerlines), means (triangles), 1st and 3rd quartiles (hinges), and minima and maxima (whiskers).

Such triple-omics integration also sheds light on the quantitative contributions of different epigenetic regulation mechanisms (Methods). Among mCH, mCG and chromatin accessibility, we found that the mCH level had the highest predictive power for gene expression in cortical neurons (average *R*^2^ = 0.188). When all epigenetic layers were considered, the expression predictability increased further (average *R*^2^ = 0.238), suggesting the presence of nonredundant contributions (Fig. 3f). Among the neurons of different layers, DNA methylation (especially mCH) exhibited slightly higher predictability for gene expression in deeper layers than in superficial layers, whereas the reverse situation held for chromatin accessibility (Supplementary Fig. 7a). Across all genes, the predictability of gene expression was generally correlated among the different epigenetic layers (Supplementary Fig. 7b). We also observed varying associations with gene characteristics. For example, mCH had higher expression predictability for longer genes, which was consistent with previous studies^17, 38^, while chromatin accessibility contributed more to genes with higher expression variability (Supplementary Fig. 7c).

### Model-based regulatory inference with GLUE

The incorporation of a graph explicitly modeling regulatory interactions in GLUE further enables a Bayesian-like approach that combines prior knowledge and observed data for posterior regulatory inference. Specifically, since the feature embeddings are designed to reconstruct the knowledge-based guidance graph and single-cell multi-omics data simultaneously (Fig. 1), their cosine similarities should reflect information from both aspects, which we adopt as “regulatory scores”.

As a demonstration, we employed the official PBMC (peripheral blood mononuclear cell) Multiome dataset from 10x^31^ and fed it to GLUE as unpaired scRNA-seq and scATAC-seq data. To capture remote cis-regulatory interactions, we employed a long-range guidance graph connecting ATAC peaks and RNA genes within 150 kb windows weighted by a power-law function that models chromatin contact probability^39, 40^ (Methods). Visualization of cell embeddings confirmed that the GLUE alignment was correct and accurate (Supplementary Fig. 8a, b). As expected, we found that the regulatory score was negatively correlated with genomic distance (Fig. 4a) and positively correlated with the empirical peak-gene correlation (computed with paired cells, Fig. 4b), with robustness across different random seeds (Supplementary Fig. 8c).

**Fig. 4.**
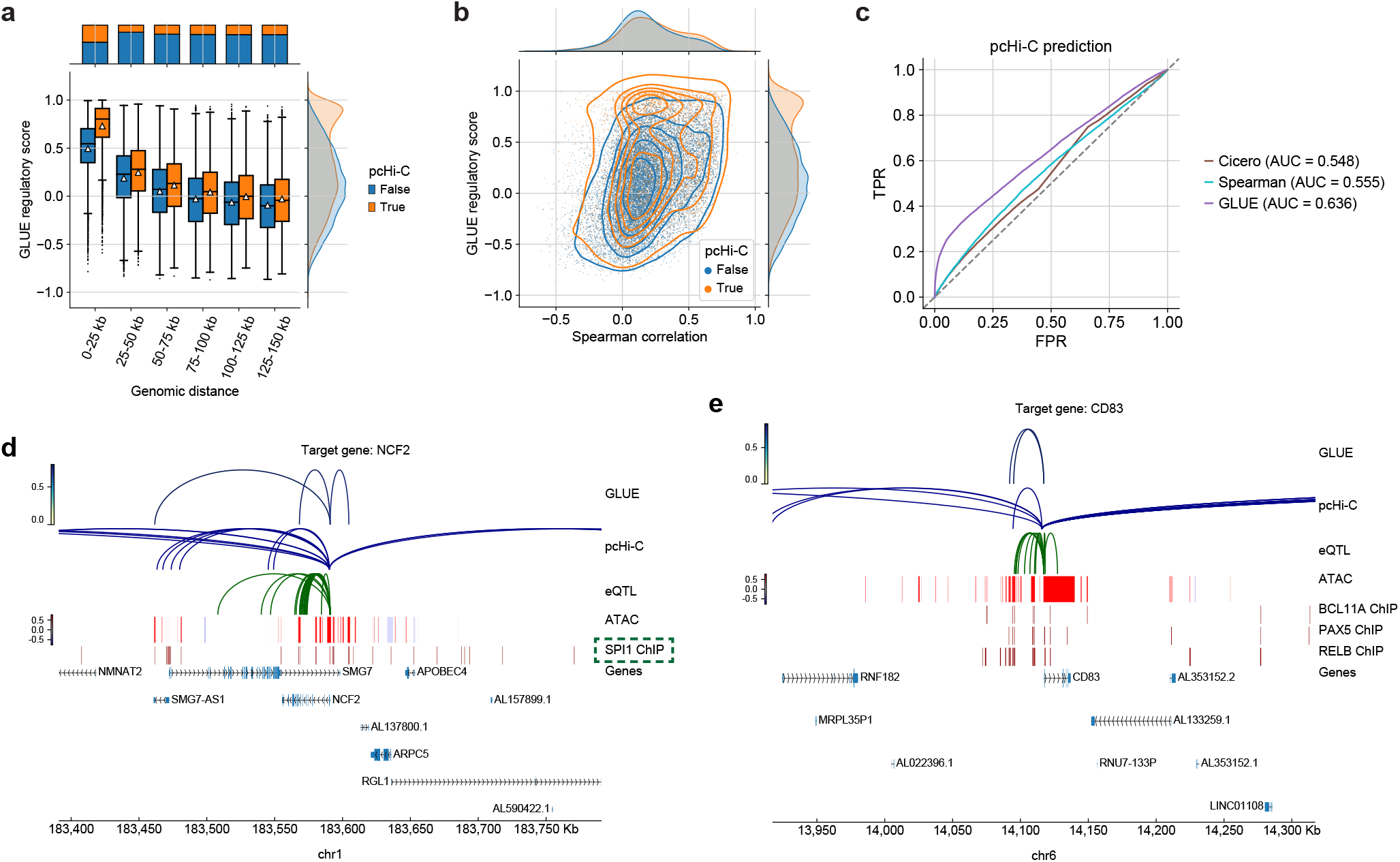
Model-based regulatory inference in PBMC. **a**, GLUE regulatory scores for peak-gene pairs across different genomic ranges, grouped by whether they had pcHi-C support. The box plots indicate the medians (centerlines), means (triangles), 1st and 3rd quartiles (hinges), and minima and maxima (whiskers). **b**, Comparison between the GLUE regulatory scores and the empirical peak-gene correlations computed on paired cells. Peak-gene pairs are colored by whether they had pcHi-C support. **c**, ROC (receiver operating characteristic) curves for predicting pcHi-C interactions based on different peak-gene association scores. **d, e**, GLUE-identified cis-regulatory interactions for **d**, *NCF2*, and **e**, *CD83*, along with individual regulatory evidence. SPI1 (highlighted with a green box) is a known regulator of *NCF2*.

To further assess whether the score reflected actual cis-regulatory interactions, we compared it with external evidence, including pcHi-C^41^ and eQTL^42^. The GLUE regulatory score was higher for pcHi-C-supported peak-gene pairs in all distance ranges (Fig. 4a) and was a better predictor of pcHi-C interactions than empirical peak-gene correlations (Fig. 4b, c), as well as Cicero^40^, the coaccessibility-based regulatory prediction method (Fig. 4c). The same held for eQTL (Supplementary Fig. 8d-f).

The GLUE framework also allows additional regulatory evidence, such as pcHi-C, to be incorporated intuitively via the guidance graph. Thus, we further trained new models with a composite guidance graph containing distance-weighted interactions as well as pcHi-C- and eQTL-supported interactions (Supplementary Fig. 9). While the multi-omics alignment was insensitive to these changes, the GLUE-derived TF-target gene network (Methods) showed more significant agreement with manually curated connections in the TRRUST v2 database^43^ than individual evidence-based networks (Supplementary Fig. 9e, Supplementary Fig. 10, Supplementary Table 3).

Interestingly, we noticed that the GLUE-inferred cis-regulatory interactions could provide new hints about the regulatory mechanisms of known TF-target pairs. For example, SPI1 is a known regulator of the *NCF2* gene, and both are highly expressed in monocytes (Supplementary Fig. 11a, b). GLUE identified three remote regulatory peaks for *NCF2* with various pieces of evidence, i.e., ∼120 kb downstream, ∼25 kb downstream, and ∼20 kb upstream from the TSS (transcription start site) (Fig. 4d), all of which were bound by SPI1. Meanwhile, most putative regulatory interactions were previously unknown. For example, *CD83* was linked with two regulatory peaks (∼25 kb upstream from the TSS), which were enriched for the binding of three TFs (BCL11A, PAX5, and RELB; Fig. 4e). While *CD83* was highly expressed in both monocytes and B cells, the inferred TFs showed more constrained expression patterns (Supplementary Fig. 11c-f), suggesting that its active regulators might differ per cell type. Supplementary Fig. 12 shows more examples of GLUE-inferred regulatory interactions.

### Atlas-scale integration over millions of cells with GLUE

As technologies continue to evolve, the throughput of single-cell experiments is constantly increasing. Recent studies have generated human cell atlases for gene expression^25^ and chromatin accessibility^26^ containing millions of cells. The integration of these atlases poses a significant challenge to computational methods due to the sheer volume of data, extensive heterogeneity, low coverage per cell, and unbalanced cell type compositions, and has yet to be accomplished at the single-cell level.

Implemented as parametric neural networks with minibatch optimization, GLUE delivers superior scalability with a sublinear time cost, promising its applicability at the atlas scale (Supplementary Fig. 13a). Using an efficient multistage training strategy for GLUE (Methods), we successfully integrated the gene expression and chromatin accessibility data into a unified multi-omics human cell atlas (Fig. 5).

**Fig. 5.**
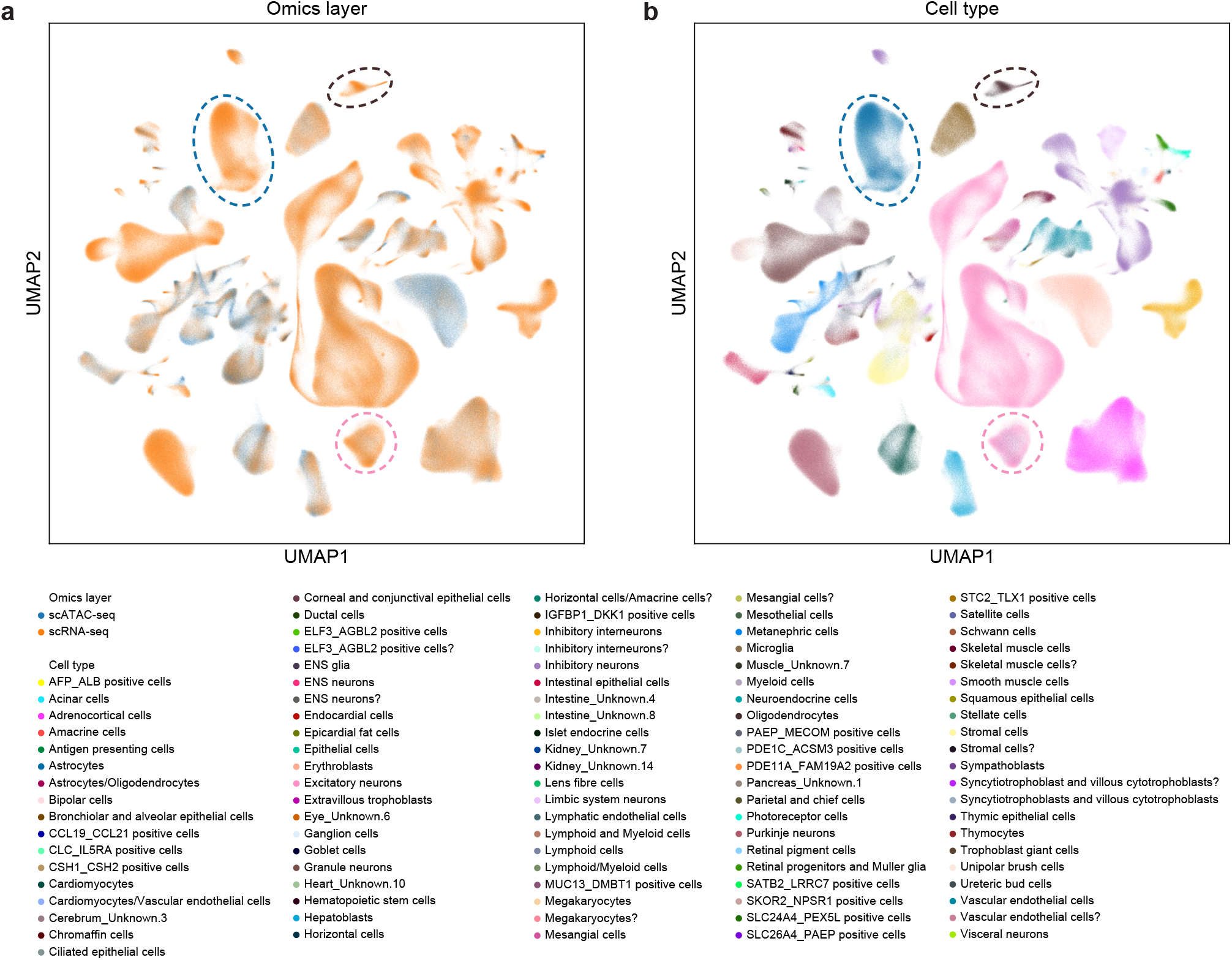
Integration of a multi-omics human cell atlas. UMAP visualizations of the integrated cell embeddings, colored by **a**, omics layers, and **b**, cell types. The pink circles highlight cells labeled as “Excitatory neurons” in scRNA-seq but “Astrocytes” in scATAC-seq. The blue circles highlight cells labeled as “Astrocytes” in scRNA-seq but “Astrocytes/Oligodendrocytes” in scATAC-seq. The brown circles highlight cells labeled as “Oligodendrocytes” in scRNA-seq but “Astrocytes/Oligodendrocytes” in scATAC-seq.

While the aligned atlas was largely consistent with the original annotations^26^ (Supplementary Fig. 13c-e), we also noticed several discrepancies. For example, cells originally annotated as “Astrocytes” in scATAC-seq were aligned to an “Excitatory neurons” cluster in scRNA-seq (highlighted with pink circles/flows in Supplementary Fig. 13). Further inspection revealed that canonical radial glial (RG) markers such as *PAX6, HES1*, and *HOPX*^44, 45^ were actively transcribed in this cluster, both in the RNA and ATAC domain (Supplementary Fig. 14), with chromatin priming^9^ also detected at both neuronal and glial markers (Supplementary Fig. 15-17), suggesting that the cluster consists of multipotent neural progenitors (likely RGs) rather than excitatory neurons or astrocytes as originally annotated. GLUE-based integration also resolved several scATAC-seq clusters that were ambiguously annotated. For example, the “Astrocytes/Oligodendrocytes” cluster was split into two halves and aligned to the “Astrocytes” and “Oligodendrocytes” clusters of scRNA-seq (highlighted with blue and brown circles/flows in Supplementary Fig. 13, respectively), which was also supported by marker expression and accessibility (Supplementary Fig. 16, 17). These results demonstrate the unique value of atlas-scale multi-omics integration.

## Discussion

Combining omics-specific autoencoders with graph-based coupling and adversarial alignment, we designed and implemented the GLUE framework for unpaired single-cell multi-omics data integration with superior accuracy and robustness. By modeling regulatory interactions across omics layers explicitly, GLUE uniquely supports model-based regulatory inference for unpaired multi-omics datasets, exhibiting even higher reliability than regular correlation analysis on paired datasets (notably, in a Bayesian interpretation, the GLUE regulatory inference can be seen as a posterior estimate, which can be continuously refined upon the arrival of new data). Furthermore, benefitting from a neural network-based design, GLUE enables notable scalability for whole-atlas alignment over millions of unpaired cells, which remains a serious challenge for *in silico* integration. In fact, we also attempted to perform integration using online iNMF, which was the only other method capable of integrating the data at full scale, but the result was far from optimal (Supplementary Fig. 18a, b). Meanwhile, an attempt to integrate the data as aggregated metacells (Methods) via the popular Seurat v3 method also failed (Supplementary Fig. 18c, d).

Unpaired multi-omics integration, also referred to as diagonal integration^14^, shares some conceptual similarities with batch effect correction^46^, as both call for the alignment of unpaired cells in certain data representations. Nonetheless, the former is significantly more challenging because of the distinct, omics-specific feature spaces. While completely unsupervised multi-omics integration has been proposed^21, 22^, such an approach is exceedingly difficult and has largely been limited to aligning continuous trajectories. For general-case multi-omics integration, additional prior knowledge is necessary. At the omics feature level, presumed feature interactions have been used via feature conversion^15-17, 30^ or coupled matrix factorization^19, 20^. While feature conversion may seem to be a straightforward solution, the inevitable information loss^18^ can have a detrimental effect on performance. Apart from the feature-converted data, Seurat v3^15^ and bindSC^30^ also devised heuristic strategies to utilize information in the original feature space, which probably explains their improved performance than methods that do not^16, 17^. At the cell level, known cell types have also been used via (semi-)supervised learning^47, 48^, but this approach incurs substantial limitations in terms of applicability since such supervision is typically unavailable and in many cases serves as the purpose of multi-omics integration *per se*^26^. Notably, one of these methods was proposed with a similar autoencoder architecture and adversarial alignment^48^, but it relied on matched cell types or clusters to orient the alignment. In fact, GLUE shares more conceptual similarity with the coupled matrix factorization methods, but with superior accuracy, robustness and scalability, which mostly benefits from its deep generative model-based design.

We note that the current framework also works for integrating omics layers with shared features, by using either the same vertex or connected surrogate vertices for each shared feature in the guidance graph. In particular, the integration between scRNA-seq and spatial transcriptomics^49, 50^ could be naturally implemented in this way. After the integration, genes not detected in the spatial transcriptome could be further imputed via cross-layer translation, through a combination of the spatial transcriptomics encoder and the scRNA-seq decoder.

As a generalizable framework, GLUE features a modular design, where the data and graph autoencoders are independently configurable.

- The data autoencoders in GLUE are customizable with appropriate generative models that conform to omics-specific data distributions. In the current work, we used the negative binomial distribution for scRNA-seq and scATAC-seq, and the zero-inflated log-normal distribution for snmC-seq (Methods). Nevertheless, generative distributions can be easily reconfigured to accommodate other omics layers, such as protein abundance^51^ and histone modification^52^, and to adopt new advances in data modeling techniques^53^.
- The guidance graphs used in GLUE have currently been limited to multipartite graphs, containing only edges between features of different layers. Nonetheless, graphs, as intuitive and flexible representations of regulatory knowledge, can embody more complex regulatory patterns, including within-modality interactions, non-feature vertices, and multi-relations. Beyond canonical graph convolution, more advanced graph neural network architectures^54-56^ may also be adopted to extract richer information from the regulatory graph.

Recent advances in experimental multi-omics technologies have increased the availability of paired data^8-11, 31^. While most of the current simultaneous multi-omics protocols still suffer from lower data quality or throughput than that of single-omics methods^57^, paired cells can be highly informative in anchoring different omics layers and should be utilized in conjunction with unpaired cells whenever available. It is straightforward to extend the GLUE framework to incorporate such pairing information, e.g., by adding another loss term that penalizes the embedding distances between paired cells^58^. Such an extension may ultimately lead to a solution for the general case of mosaic integration^14^.

Apart from multi-omics integration, we also note that the GLUE framework could be suitable for cross-species integration, especially when distal species are concerned and one-to-one orthologs are limited. Specifically, we may compile all orthologs into a GLUE guidance graph and perform integration without explicit ortholog conversion. Under that setting, the GLUE approach could also be conceptually connected to a recent work called SAMap^59^.

We believe that GLUE, as a modular and generalizable framework, creates an unprecedented opportunity towards effectively delineating gene regulatory maps via large-scale multi-omics integration at single-cell resolution. The whole package of GLUE, along with tutorials and demo cases, is available online at https://github.com/gao-lab/GLUE for the community.

## Methods

### The GLUE framework

We assume that there are *K* different omics layers to be integrated, each with a distinct feature set 𝒱_*k*_, *k* = 1,2, …, *K*. For example, in scRNA-seq, 𝒱_*k*_ is the set of genes, while in scATAC-seq, 𝒱_*k*_ is the set of chromatin regions. The data spaces of different omics layers are denoted as 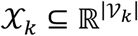 with varying dimensionalities. We use **x**_*k*_^(*n*)^ ∈ 𝒳_*k*_, *n* = 1,2, …, *N*_*K*_ to denote cells from the *k*^th^ omics layer and 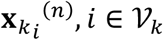 to denote the observed value of feature *i* of the *k*^th^ layer in the *n*^th^ cell. *N*_*K*_ is the sample size of the *k*^th^ layer. Notably, the cells from different omics layers are unpaired and can have different sample sizes. To avoid cluttering, we drop the superscript (*n*) when referring to an arbitrary cell.

We model the observed data from different omics layers as generated by a low-dimensional latent variable (i.e., cell embedding) **u** ∈ ℝ^*m*^:

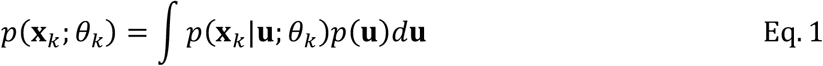

where *p*(**u**) is the prior distribution of the latent variable, *p*(**x**_*k*_|**u**; *θ*_*k*_) are learnable generative distributions (i.e., data decoders), and *θ*_*k*_ denotes learnable parameters in the decoders. The cell latent variable **u** is shared across different omics layers. In other words, **u** represents the common cell states underlying all omics observations, while the observed data from each layer are generated by a specific type of measurement of the underlying cell states.

With the introduction of variational posteriors *q*(**u**|**x**_*k*_; *ϕ*_*k*_) (i.e., data encoders, where *ϕ*_*k*_ are learnable parameters in the encoders), model fitting can be efficiently performed by maximizing the following evidence lower bounds:

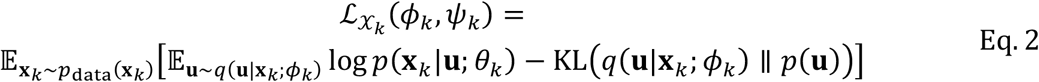

Since different autoencoders are independently parameterized and trained on separate data, the cell embeddings learned for different omics layers could have inconsistent semantic meanings unless they are linked properly.

To link the autoencoders, we propose a guidance graph 𝒢 = (𝒱, ℰ), which incorporates prior knowledge about the regulatory interactions among features at distinct omics layers, where 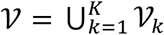 is the universal feature set and ℰ = {(*i, j*)|*i, j* ∈ 𝒱} is the set of edges. Each edge is also associated with signs and weights, which are denoted as *s*_*ij*_ and *w*_*ij*_, respectively. We require that *w*_*ij*_ ∈ (0,1], which can be interpreted as interaction credibility, and that *s*_*ij*_ ∈ {−1,1}, which specifies the sign of the regulatory interaction. For example, an ATAC peak located near the promoter of a gene is usually assumed to positively regulate its expression, so they can be connected with a positive edge (*s*_*ij*_ = 1). Meanwhile, DNA methylation in the gene promoter is usually assumed to suppress expression, so they can be connected with a negative edge (*s*_*ij*_ = −1). In addition to the connections between features, self-loops are also added for numerical stability, with *s*_*ii*_ = 1, *w*_*ii*_ = 1, ∀*i* ∈ 𝒱.

We treat the guidance graph as observed variable and model it as generated by low-dimensional feature latent variables (i.e., feature embeddings) **v**_*i*_ ∈ ℝ^*m*^, *i* ∈ 𝒱. Furthermore, differing from the previous model, we now model **x**_*k*_ as generated by the combination of feature latent variables **v**_*i*_ ∈ ℝ^*m*^, *i* ∈ 𝒱_*k*_ and the cell latent variable **u** ∈ ℝ^*m*^. For convenience, we introduce the notation **V** ∈ ℝ^*m*×|𝒱|^, which combines all feature embeddings into a single matrix. The model likelihood can thus be written as:

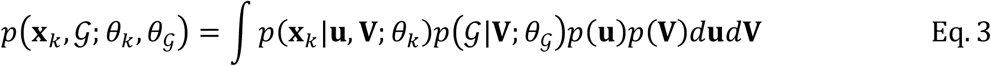

where *p*(**x**_*k*_|**u, V**; *θ*_*k*_) and *p*(𝒢|**V**; *θ*_𝒢_) are learnable generative distributions for the omics data (i.e., data decoders) and knowledge graph (i.e., graph decoder), respectively. *θ*_*k*_ and *θ*_𝒢_ are learnable parameters in the decoders. *p*(**u**) and *p*(**V**) are the prior distributions of the cell latent variable and feature latent variables, respectively, which are fixed as standard normal distributions for simplicity:

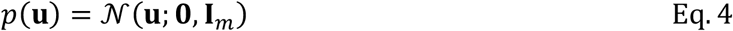

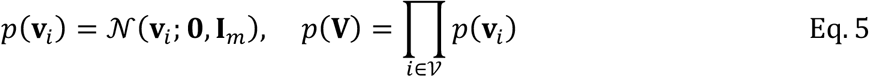

though alternatives may also be used^60^. For convenience, we also introduce the notation 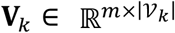, which contains only feature embeddings in the *k*^th^ omics layer, and **u**_*k*_, which emphasizes that the cell embedding is from a cell in the *k*^th^ omics layer.

The graph likelihood *p*(𝒢|**V**; *θ*_𝒢_) (i.e., graph decoder) is defined as:

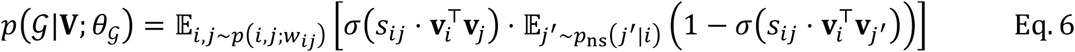

where *σ* is the sigmoid function and *p*_*ns*_ is a negative sampling distribution^61^. In other words, we first sample the edges (*i, j*) with probabilities proportional to the edge weights and then sample vertices *j*’ that are not connected to *i* and treat them as if *s*_*ij*′_ = *s*_*ij*_. When maximizing the graph likelihood, the inner products between features are maximized or minimized (per edge sign) based on the Bernoulli distribution. For example, ATAC peaks located near the promoter of a gene would be encouraged to have similar embeddings to that of the gene, while DNA methylation in the gene promoter would be encouraged to have a dissimilar embedding to that of the gene.

The data likelihoods *p*(**x**_*k*_|**u, V**; *θ*_*k*_) (i.e., data decoders) in Eq. 3 are built upon the inner product between the cell embedding **u** and feature embeddings **V**_*k*_. Thus, analogous to the loading matrix in principal component analysis (PCA), the feature embeddings **V**_*k*_ confer semantic meanings for the cell embedding space. As **V**_*k*_ are modulated by interactions among omics features in the guidance graph, the semantic meanings become linked. The exact formulation of data likelihood depends on the omics data distribution. For example, for count-based scRNA-seq and scATAC-seq data, we used the negative binomial (NB) distribution:

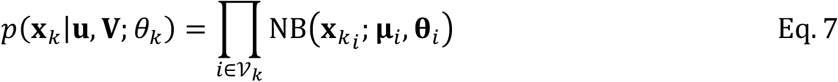

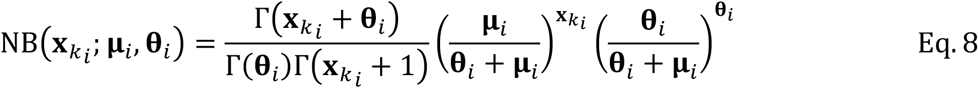

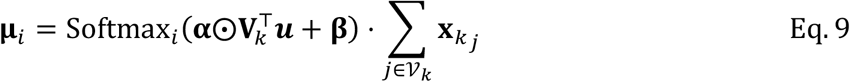

where 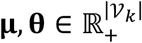 are the mean and dispersion of the NB distribution, respectively, and 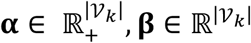 are scaling and bias factors. ⨀ is the Hadamard product. Softmax_*i*_ represents the *i*^th^ dimension of the softmax output. The set of learnable parameters is *θ*_*k*_ = {**θ, α, β**}. Analogously, many other distributions can also be supported, as long as we can parameterize the means of the distributions by feature-cell inner products.

For efficient inference and optimization, we introduce the following factorized variational posterior:

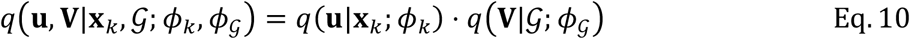

The graph variational posterior *q*(**V**|𝒢; *ϕ*_𝒢_) (i.e., graph encoder) is modeled as diagonal-covariance normal distributions parameterized by a graph convolutional network^62^:

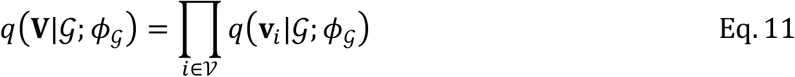

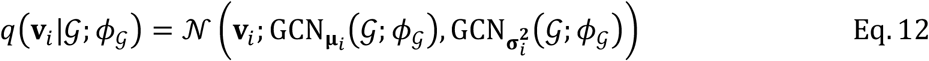

where *ϕ*_𝒢_ represents the learnable parameters in the GCN encoder.

The variational data posteriors *q*(**u**|**x**_*k*_; *ϕ*_*k*_) (i.e., data encoders) are modeled as diagonal-covariance normal distributions parameterized by multilayer perceptron (MLP) neural networks:

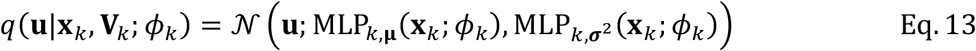

where *ϕ*_*k*_ is the set of learnable parameters in the MLP encoder of the *k*^th^ omics layer.

Model fitting can then be performed by maximizing the following evidence lower bound:

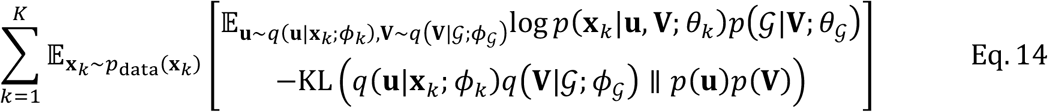

which can be further rearranged into the following form:

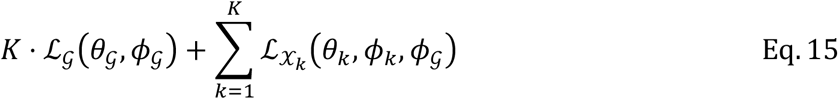

where we have

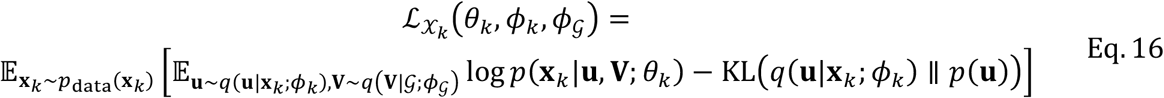

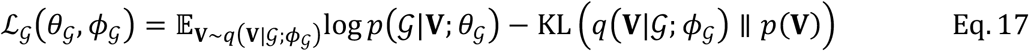

Below, for convenience, we denote the union of all encoder parameters as 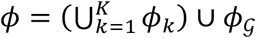 and the union of all decoder parameters as 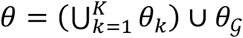.

To ensure the proper alignment of different omics layers, we use the adversarial alignment strategy^28,63^. A discriminator D with a *K*-dimensional softmax output is introduced, which predicts the omics layers of cells based on their embeddings **u**. The discriminator D is trained by minimizing the multiclass classification cross entropy:

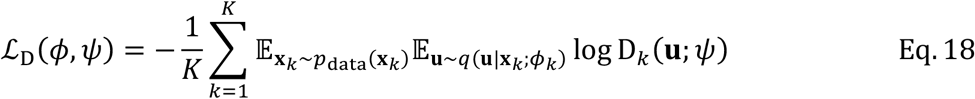

where D_*k*_ represents the *k*^th^ dimension of the discriminator output and *ψ* is the set of learnable parameters in the discriminator. The data encoders can then be trained in the opposite direction to fool the discriminator, ultimately leading to the alignment of cell embeddings from different omics layers^64^.

The overall training objective of GLUE thus consists of:

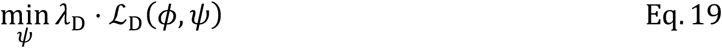

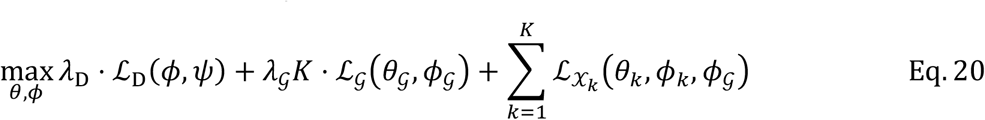

The two hyperparameters *λ*_D_ and *λ*_𝒢_ control the contributions of adversarial alignment and graph-based feature embedding, respectively. We use stochastic gradient descent (SGD) to train the GLUE model. Each SGD iteration is divided into two steps. In the first step, the discriminator is updated according to objective Eq. 19. In the second step, the data and graph autoencoders are updated according to Eq. 20. The RMSprop optimizer with no momentum term is employed to ensure the stability of adversarial training.

### Implementation details

We applied linear dimensionality reduction using canonical methods such as PCA (for scRNA-seq) or LSI (latent semantic indexing, for scATAC-seq) as the first transformation layers of the data encoders (note that the decoders were still fitted in the original feature spaces). This effectively reduced model size and enabled a modular input, so advanced dimensionality reduction or batch effect correction methods can also be used instead as preprocessing steps for GLUE integration.

To ensure stable alignment, we used batch normalization in the data encoder layers and employed additive noise annealing. Specifically, noise **ϵ** ∼ 𝒩(**ϵ**; **0**, *τ* ⋅ **I**_*m*_) was added to the cell embeddings **u** before passing to the discriminator. The parameter *τ* controls the noise level, which starts at *τ* = 1 and decreases linearly per epoch until reaching 0 (i.e., noise annealing). The number of annealing epochs was set automatically based on the data size and learning rate to match a learning progress equivalent to 4,000 iterations at a learning rate of 0.002.

During model training, 10% of the cells were used as the validation set. In the final stage of training, the learning rate would be reduced by factors of 10 if the validation loss did not improve for consecutive epochs. Training would be terminated if the validation loss still did not improve for consecutive epochs. The patience for learning rate reduction, training termination, and the maximal number of training epochs were automatically set based on the data size and learning rate to match a learning progress equivalent to 1,000, 2,000, and 16,000 iterations at a learning rate of 0.002, respectively.

For all benchmarks and case studies with GLUE, we used the default hyperparameters unless explicitly stated. The set of default hyperparameters is presented in Supplementary Fig. 4.

### Systematic benchmarks

UnionCom^21^ and GLUE were executed using the Python packages “unioncom” (v0.3.0) and “scglue” (v0.1.1), respectively. Online iNMF^16^, LIGER^17^, bindSC^30^, and Seurat v3^15^ were executed using the R packages “rliger” (v1.0.0), “rliger” (v1.0.0), “bindSC” (v1.0.0), and “Seurat” (v4.0.2), respectively. For each method, we used the default hyperparameter settings and data preprocessing steps as recommended. For the scRNA-seq data, 2,000 highly variable genes were selected using the Seurat “vst” method. To construct the guidance graph, we connected ATAC peaks with RNA genes via positive edges if they overlapped in either the gene body or proximal promoter regions (defined as 2 kb upstream from the TSS). For the methods that require feature conversion (online iNMF, LIGER, bindSC, and Seurat v3), we converted the scATAC-seq data to gene-level activity scores by summing up counts in the ATAC peaks connected to specific genes in the guidance graph. Notably, online iNMF and LIGER also recommend an alternative way of ATAC feature conversion, i.e., directly counting ATAC fragments falling in gene body and promoter regions without resorting to ATAC peaks (https://htmlpreview.github.io/?https://github.com/welch-lab/liger/blob/master/vignettes/Integrating_scRNA_and_scATAC_data.html), which we abbreviate as FiG (fragments in genes). We also tested the FiG feature conversion method with online iNMF and LIGER.

MAP (mean average precision) was used to evaluate the cell type resolution. Supposing that the cell type of the *i*^th^ cell is *y*^(*i*)^ and that the cell types of its *K* ordered nearest neighbors are 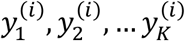, the MAP is then defined as follows:

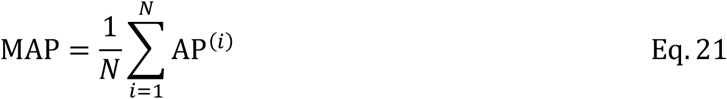

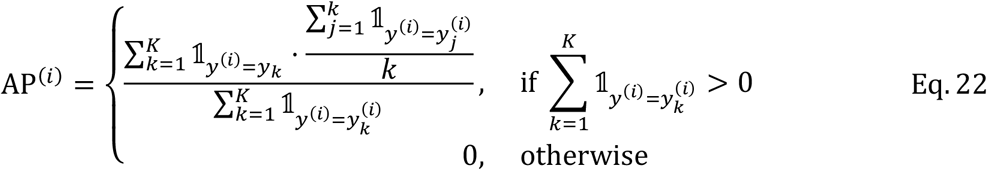

where 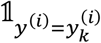 is an indicator function that equals 1 if 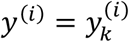 and 0 otherwise. For each cell, AP (average precision) computes the average cell type precision up to each cell type-matched neighbor, and MAP is the average AP across all cells. We set *K* to 1% of the total number of cells in each dataset. MAP has a range of 0 − 1, and higher values indicate better cell type resolution.

SAS (Seurat alignment score) was used to evaluate the extent of mixing among distinct omics layers and was computed as described in the original paper^32^:

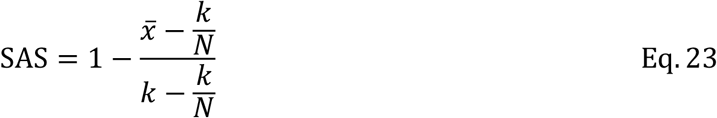

where 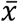 is the average number of cells from the same omics layer among the *k* nearest neighbors (different layers were first subsampled to the same number of cells as the smallest layer), and *N* is the number of omics layers. We set *k* to 1% of the subsampled cell number. SAS has a range of 0 − 1, and higher values indicate better mixing.

FOSCTTM (fraction of samples closer than the true match)^33^ was used to evaluate the single-cell level alignment accuracy. It was computed on two datasets with known cell-to-cell pairings. Suppose that each dataset contains *N* cells, and that the cells are sorted in the same order, i.e., the *i*^th^ cell in the first dataset is paired with the *i*^th^ cell in the second dataset. Denote **x** and **y** as the cell embeddings of the first and second dataset, respectively. The FOSCTTM is then defined as:

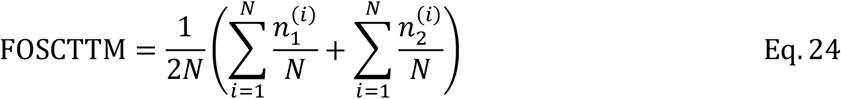

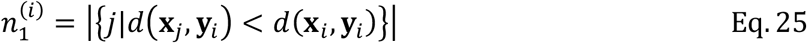

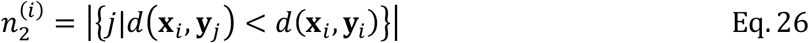

where 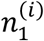 and 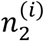 are the number of cells in the first and second dataset, respectively, that are closer to the *i*^th^ cell than their true matches in the opposite dataset. *d* is the Euclidean distance.

For the baseline benchmark, each method was run 8 times with different random seeds, except for bindSC, which has a deterministic implementation and was run only once. For the guidance corruption benchmark, we removed the specified proportions of existing peak-gene interactions and added equal numbers of nonexistent interactions, so the total number of interactions remained unchanged. Of note, feature conversion was also repeated using the corrupted guidance graphs. The corruption procedure was repeated 8 times with different random seeds. For the subsampling benchmark, the scRNA-seq and scATAC-seq cells were subsampled in pairs (so FOSCTTM could still be computed). The subsampling process was also repeated 8 times with different random seeds.

For the systematic scalability test (Supplementary Fig. 13a), all methods were run on a Linux workstation with 40 CPU cores (two Intel Xeon Silver 4210 chips), 250 GB of RAM, and NVIDIA GeForce RTX 2080 Ti GPUs. Only a single GPU card was used when training GLUE.

### Triple-omics integration

The scRNA-seq and scATAC-seq data were handled as previously described (see the section “Systematic benchmarks”). Due to low coverage per single-C site, the snmC-seq data were converted to average methylation levels in gene bodies. The mCH and mCG levels were quantified separately, resulting in 2 features per gene. The gene methylation levels were normalized by the global methylation level per cell. An initial dimensionality reduction was performed using PCA (see the section “Implementation details”). For the triple-omics guidance graph, the mCH and mCG levels were connected to the corresponding genes with negative edges.

The normalized methylation levels were positive, with dropouts corresponding to the genes that were not covered in single cells. As such, we used the zero-inflated log-normal (ZILN) distribution for the data decoder:

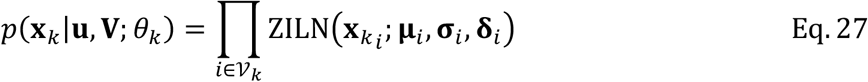

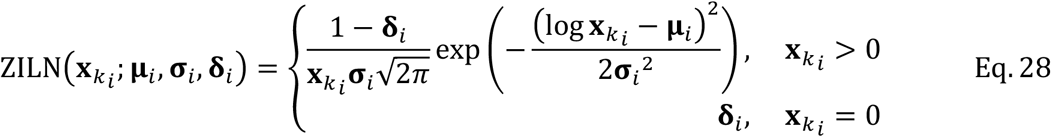

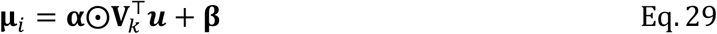

where 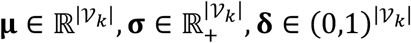 are the log-scale mean, log-scale standard deviation and zero-inflation parameters of the ZILN distribution, respectively, and 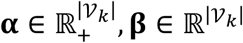 are scaling and bias factors.

To unify the cell type labels, we performed a nearest neighbor-based label transfer with the snmC-seq dataset as a reference. The 5 nearest neighbors in snmC-seq were identified for each scRNA-seq and scATAC-seq cell in the aligned embedding space, and majority voting was used to determine the transferred label. To verify whether the alignment was correct, we tested for significant overlap in cell type marker genes. The features of all omics layers were first converted to genes. Then, for each omics layer, the cell type markers were identified using the one-vs.-rest Wilcoxon rank-sum test with the following criteria: FDR < 0.05 and log-fold change > 0 for scRNA-seq/scATAC-seq; FDR < 0.05 and log-fold change < 0 for snmC-seq. The significance of marker overlap was determined by the three-way Fisher’s exact test^37^.

To perform correlation and regression analysis after the integration, we clustered all cells from the three omics layers using fine-scale k-means (k = 200). Then, for each omics layer, the cells in each cluster were aggregated into a metacell by summing their expression/accessibility counts or averaging their DNA methylation levels. Therefore, the metacells were established as paired samples, based on which feature correlation and regression analyses could be conducted.

### Model-based cis-regulatory inference

To ensure consistency of cell types, we first selected the overlapping cell types between the 10x Multiome and pcHi-C data. The remaining cell types included T cells, B cells and monocytes. The eQTL data were used as is, because they were not cell type-specific. For scRNA-seq, we selected 6,000 highly variable genes. For the initial regulatory inference, the guidance graph was constructed by connecting RNA genes with ATAC peaks within 150 kb of the gene promoters (defined as 2 kb upstream from the TSS); the graph was weighted by a power-law function *w* = (*d* + 1) ^−0.75^ (*d* is the genomic distance in kb), which has been proposed to model the probability of chromatin contact^39, 40^.

To incorporate the regulatory evidence of pcHi-C and eQTL, we anchored all evidence to that between the ATAC peaks and RNA genes. A peak-gene pair was considered supported by pcHi-C if (1) the gene promoter was within 1 kb of a bait fragment, (2) the peak was within 1 kb of an other-end fragment, and (3) significant contact was identified between the bait and the other-end fragment in pcHi-C. The pcHi-C-supported peak-gene interactions were weighted by multiplying the promoter-to-bait and the peak-to-other-end power-law weights (see above). If a peak-gene pair was supported by multiple pcHi-C contacts, the weights were summed and clipped at a maximum of 1. A peak-gene pair was considered supported by eQTL if (1) the peak overlapped an eQTL locus and (2) the locus was associated with the expression of the gene. The eQTL-supported peak-gene interactions were assigned weights of 1. The composite guidance graph was constructed by adding the pcHi-C- and eQTL-supported interactions to the previous distance-based interactions, allowing for multi-edges.

For regulatory inference, only peak-gene pairs within 150 kb in distance were considered. The GLUE training process was repeated 4 times with different random seeds. For each repeat, the peak-gene regulatory score was computed as the cosine similarity between the feature embeddings. The final regulatory inference was obtained by averaging the regulatory scores across the 4 repeats.

### TF-target gene regulatory inference

We employed the SCENIC workflow^65^ to construct a TF-gene regulatory network from the inferred peak-gene regulatory interactions. Briefly, the SCENIC workflow first constructs a gene coexpression network based on the scRNA-seq data, and then uses external cis-regulatory evidence to filter out false positives. SCENIC accepts cis-regulatory evidence in the form of gene rankings per TF, i.e., genes with higher TF enrichment levels in their regulatory regions are ranked higher. To construct the rankings based on our inferred peak-gene interactions, we first overlapped the ENCODE TF ChIP peaks^66^ with the ATAC peaks and counted the number of ChIP peaks for each TF in each ATAC peak. Since different genes can have different numbers of connected ATAC peaks, and the ATAC peaks vary in length (longer peaks can contain more ChIP peaks by chance), we devised a sampling-based approach to evaluate TF enrichment. Specifically, for each gene, we randomly sampled 1,000 sets of ATAC peaks that matched the connected ATAC peaks in both number and length distribution. We counted the numbers of TF ChIP peaks in these random ATAC peaks as null distributions. For each TF in each gene, an empirical *P* value could then be computed by comparing the observed number of ChIP peaks to the null distribution. Finally, we ranked the genes by the empirical *P* values for each TF, producing the cis-regulatory rankings used by SCENIC. Since peak-gene-based inference is mainly focused on remote regulatory regions, proximal promoters could be missed. As such, we provided SCENIC with both the above peak-based and proximal promoter-based cis-regulatory rankings.

### Integration for the human multi-omics atlas

The scRNA-seq and scATAC-seq atlases have highly unbalanced cell type compositions, which is primarily caused by differences in organ sampling sizes (Supplementary Fig. 13b). Although cell types are unknown during real-world analyses, organ sources are typically available and can be utilized to help balance the integration process. To perform organ-balanced data preprocessing, we first subsampled each omics layer to match the organ compositions. For the scRNA-seq data, 4,000 highly variable genes were selected using the organ-balanced subsample. Then, for the initial dimensionality reduction, we fitted PCA (scRNA-seq) and LSI (scATAC-seq) on the organ-balanced subsample and applied the projection to the full data. The PCA/LSI coordinates were used as the first transformation layer in the GLUE data encoders (see the section “Implementation details”), as well as for metacell aggregation (see below). The guidance graph was constructed as described previously (see the section “Systematic benchmarks”).

The two atlases consist of large numbers of cells but with low coverage per cell. To alleviate dropout and increase the training speed simultaneously, we designed a multistage training strategy, where the GLUE model was pretrained on aggregated metacells and then fine-tuned on the original single cells. Specifically, in the first stage, we clustered the cells in each omics layer using fine-scaled k-means (k = 100,000 for scRNA-seq and k = 40,000 for scATAC-seq). To balance the organ compositions at the same time, k-means centroids were fitted on the previous organ-balanced subsample and then applied to the full data. The cells in each k-means cluster were aggregated into a metacell by summing their expression/accessibility counts and averaging their PCA/LSI coordinates. GLUE was then pretrained on the aggregated metacells without adversarial alignment, which roughly oriented the cell embeddings but did not actually align them. To better utilize the large data size, the hidden layer dimensionality was doubled to 512 from the default 256.

In the second stage, GLUE was fine-tuned on the full single-cell data with weighted adversarial alignment (see below). As shown in previous work^28^, pure adversarial alignment amounts to minimizing a generalized form of Jensen–Shannon divergence among the cell embedding distributions of different omics layers:

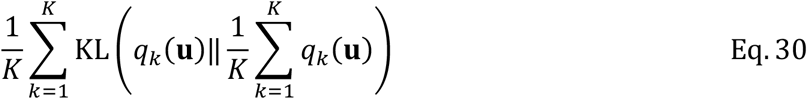

where 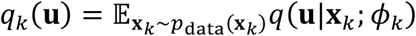 represents the marginal cell embedding distribution of the *k*^th^ layer. Without other loss terms, Eq. 30 converges at perfect alignment, i.e., when *q*_*i*_ (**u**) = *q*_*j*_ (**u**), ∀*i ≠ j*. This can be problematic when cell type compositions differ dramatically across different layers, as was the case here. To address this issue, we added cell-specific weights *w*^(*n*)^ to the discriminator loss in Eq. 18:

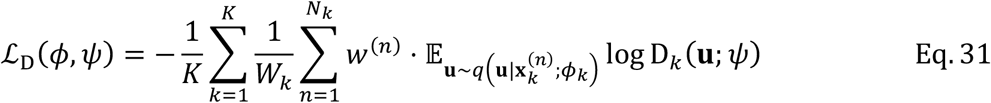

where the normalizer 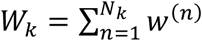. The adversarial alignment still amounts to minimizing Eq. 30 but with weighted marginal cell embedding distributions 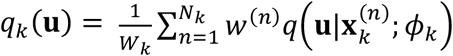. By assigning appropriate weights to balance the cell distributions across different layers, the optimum of *q*_*i*_ (**u**) = *q*_*j*_ (**u**), ∀*i* ≠ *j* could be much closer to the desired alignment. For example, a viable choice for the cell weights is organ-balancing weights. Suppose that the organ proportions in scRNA-seq and scATAC-seq are *f*_1_, *f*_2_, …, *f*_0_ and *g*_1_, *g*_2_, …, *g*_0_ (*0* is the number of organs, 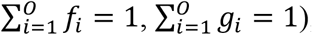), respectively. We can weight the RNA cells in the *i*^th^ organ by 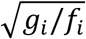 and weight the ATAC cells in the *i*^th^ organ by 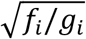, so that the *i*^th^ organ has a balanced accumulative contribution of 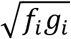, regardless of omics layers. However, in case there are major differences among the cell type compositions within the same organ, organ-level balancing can be insufficient. As such, we designed a method to compute cell type-level balancing weights in an unsupervised manner. For each omics layer, we clustered the cell embeddings using Louvain clustering and matched the clusters in different layers via cosine similarity. The population size of each cluster was distributed to its matched counterparts by cosine similarity. Subsequently, for each omics layer, we could obtain the proportion of each cluster *f*_*i*_ and its effective proportion *g*_*i*_ in the opposite layer (by normalizing the effective population size received from the opposite layer). Balancing weights could then be computed as above. The weight-balanced alignment proved effective in aligning the highly skewed data (Fig. 5).

For a comparison with other integration methods, we also tried online iNMF and Seurat v3. Online iNMF was the only other method that could scale to millions of cells, so we applied it to the full dataset. On the other hand, Seurat v3 showed the second-best accuracy in our previous benchmark. We also managed to apply it to the aggregated data as used in the first stage of GLUE training, due to the fact that Seurat v3 could not scale to the full dataset (Supplementary Fig. 13a).

## Data availability

All datasets used in this study are already published and were obtained from public data repositories. See Supplementary Table 1 for detailed information, including access codes and URLs. All benchmark data are available in Supplementary Table 2.

## Code availability

The GLUE framework was implemented in the “scglue” Python package, which is available at https://github.com/gao-lab/GLUE. For reproducibility, the scripts for all benchmarks and case studies were assembled using Snakemake, which is also available in the above repository.

## Acknowledgments

The authors would like to thank Drs. Zemin Zhang, Xiaoliang Sunney Xie, Letian Tao, Cheng Li, Jian Lu (at Peking University) and Yang Ding (at the Beijing Institute of Radiation Medicine) for their helpful discussions and comments during the study, as well as authors of the datasets used in this work for their kindly help.

This work was supported by funds from the National Key Research and Development Program (2016YFC0901603), the China 863 Program (2015AA020108), and the State Key Laboratory of Protein and Plant Gene Research and the Beijing Advanced Innovation Center for Genomics (ICG) at Peking University. The research of G.G. was supported in part by the National Program for Support of Top-notch Young Professionals.

Part of the analysis was performed on the Computing Platform of the Center for Life Sciences of Peking University and supported by the High-performance Computing Platform of Peking University.

## Author contributions

G.G. conceived the study and supervised the research; Z.J.C. designed and implemented the computational framework and conducted benchmarks and case studies, with guidance from G.G.; Z.J.C. and G.G. wrote the manuscript.

