## Supplementary Information for "Multi-omics integration and regulatory inference for unpaired single-cell data with a graph-linked unified embedding framework"

**Cao et al.**

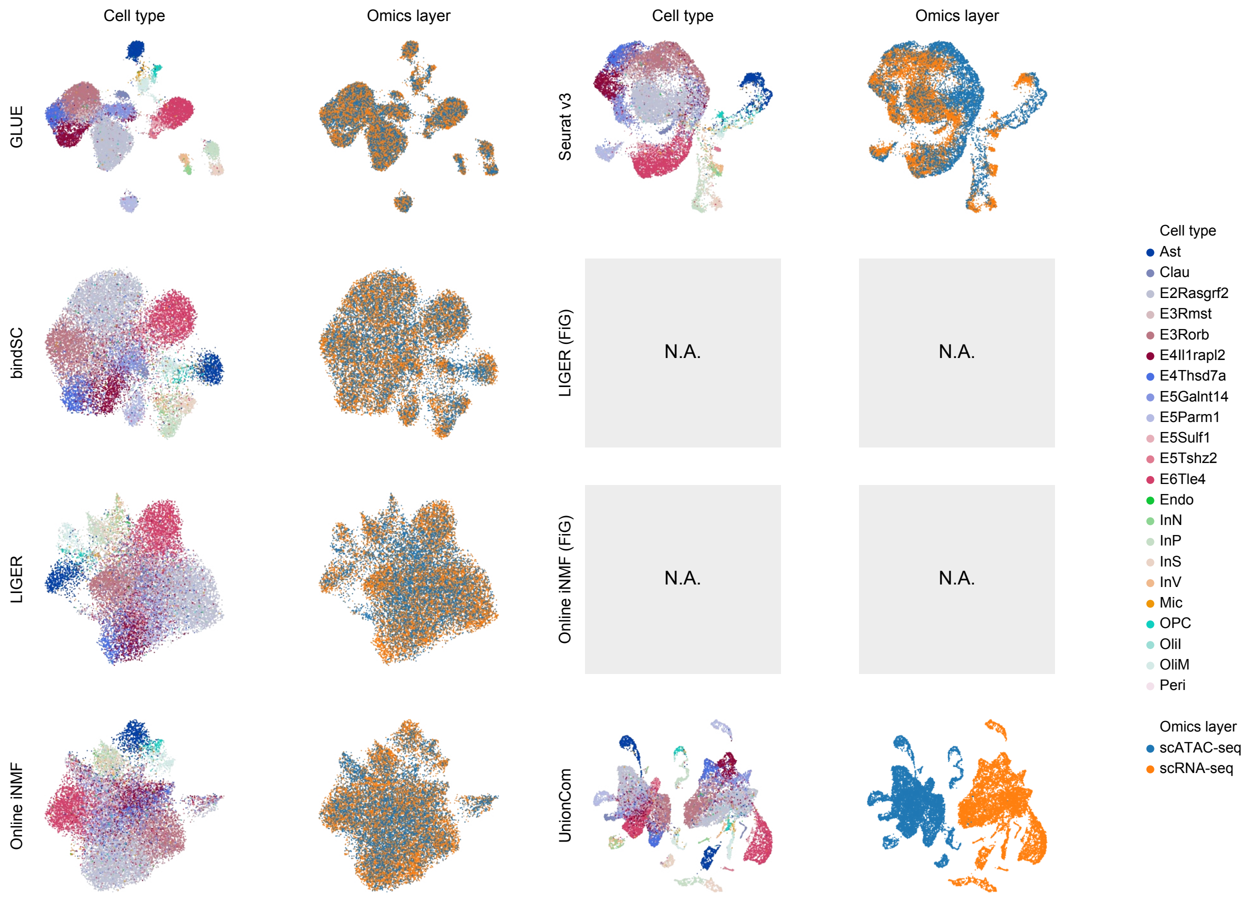


**Supplementary Fig. 1 UMAP visualizations of the cell embeddings in the SNARE-seq dataset aligned with different integration methods.**
Online iNMF and LIGER could not run with FiG conversion because raw ATAC fragment file was not available.

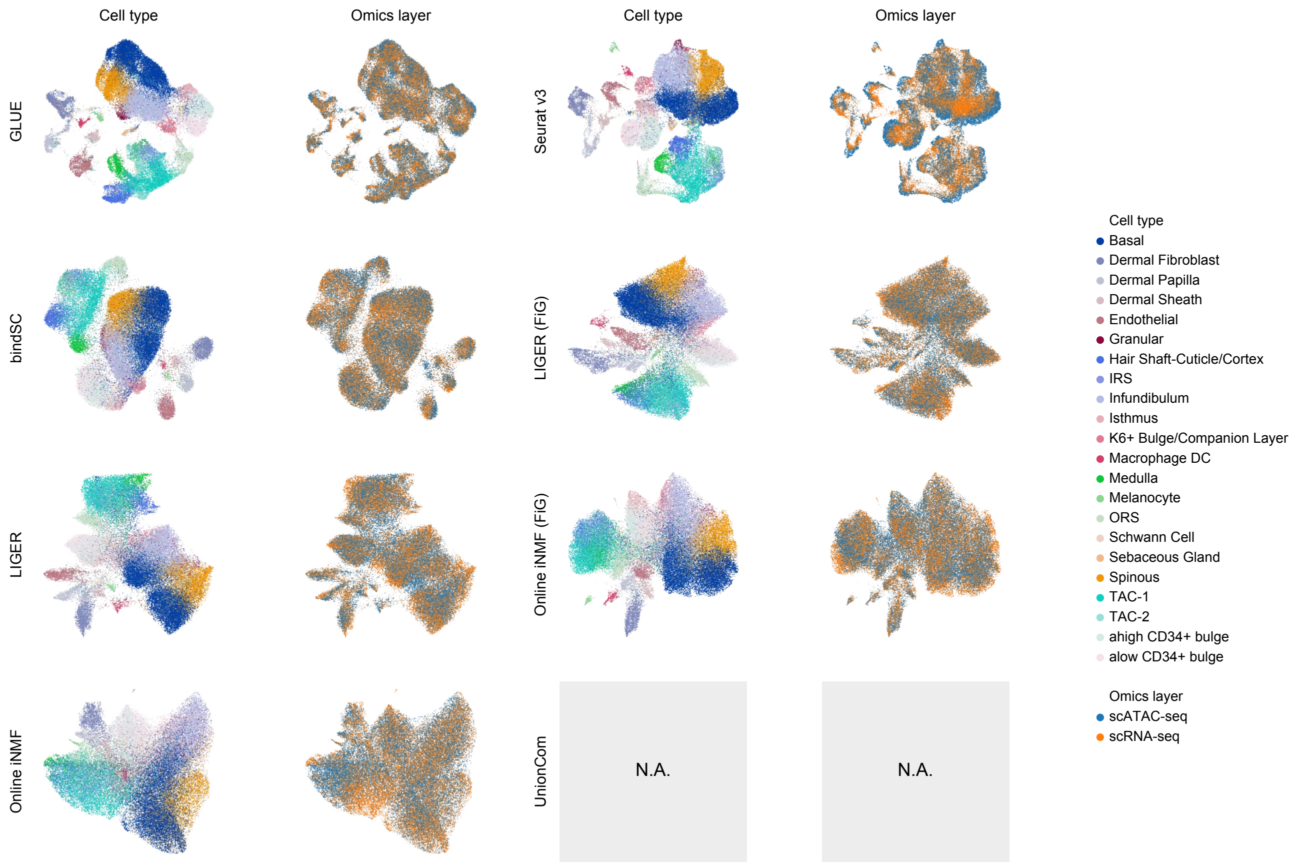


**Supplementary Fig. 2 UMAP visualizations of the cell embeddings in the SHARE-seq dataset aligned with different integration methods.**
UnionCom failed to run because of memory overflow.

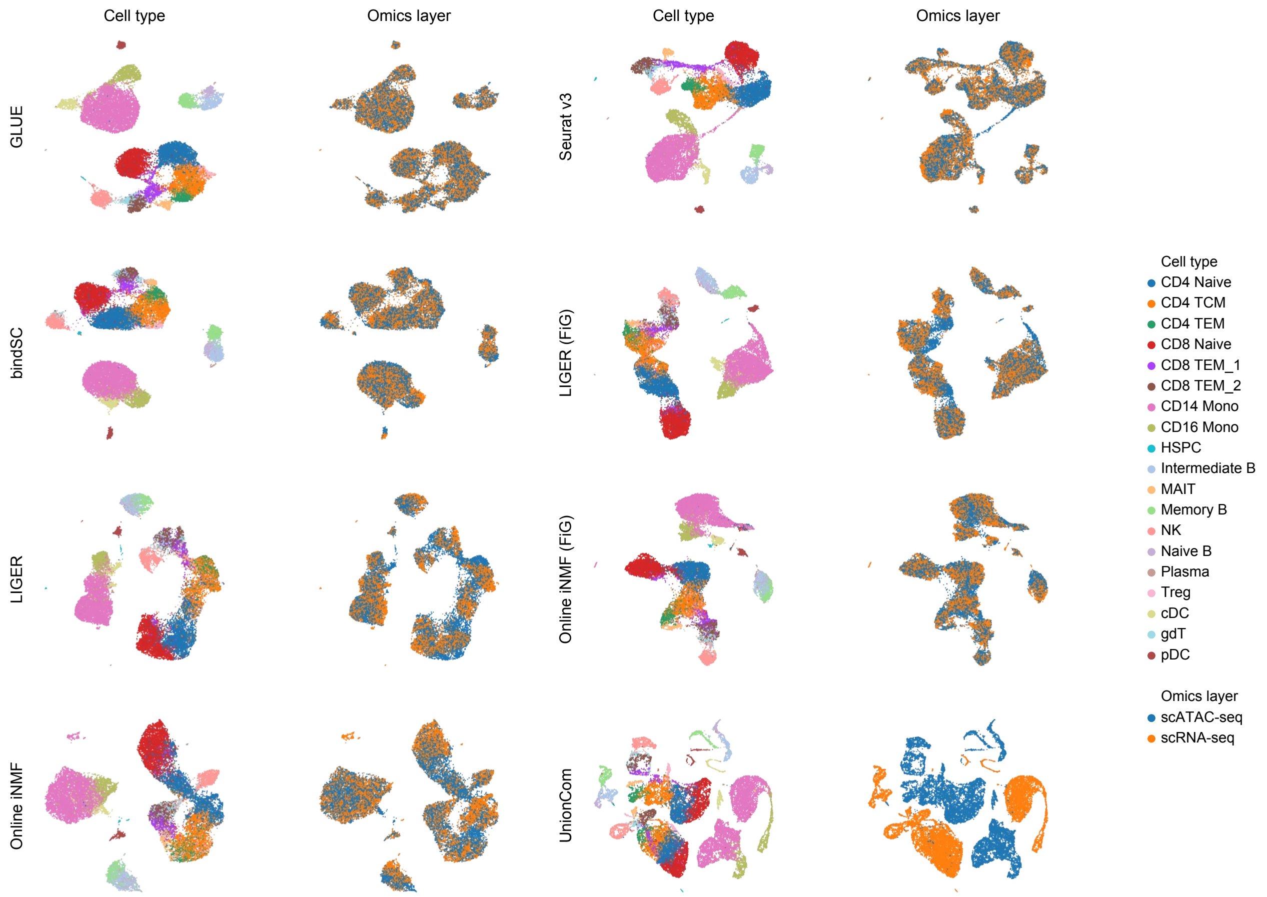


**Supplementary Fig. 3 UMAP visualizations of the cell embeddings in the 10x Multiome dataset aligned with different integration methods.**

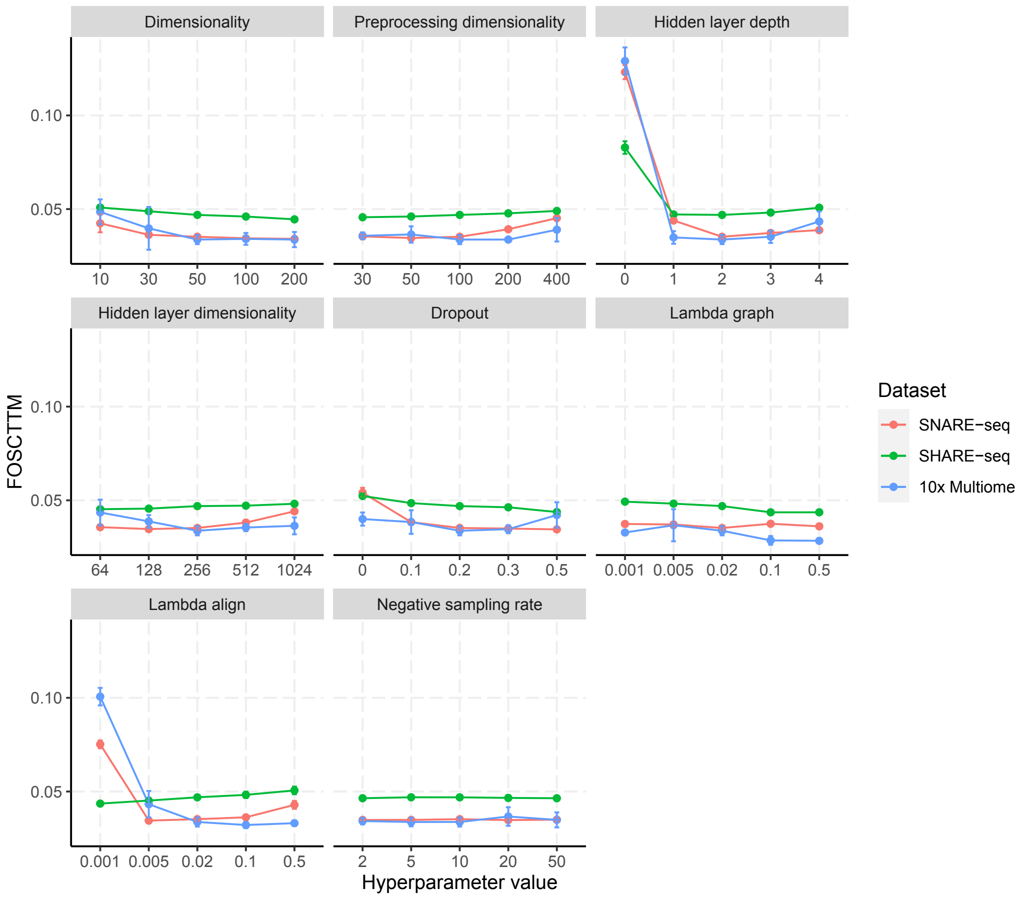


**Supplementary Fig. 4 FOSCTTM of GLUE under different hyperparameter settings.** "Dimensionality" denotes the cell embedding dimensionality. "Preprocessing dimensionality" is the reduced dimensionality used for the first transformation layers of the data encoders (see Methods). "Hidden layer depth" is the number of hidden layers in the data encoders and modality discriminator. "Hidden layer dimensionality" is the dimensionality of hidden layers in the data encoders and modality discriminator. "Dropout" is the dropout rate of hidden layers in data encoders and modality discriminator. "Lambda graph" is the weight of the graph loss ($\lambda_{\mathcal{G}}$). "Lambda align" is the weight of the adversarial alignment ($\lambda_{\mathrm{D}}$). "Negative sampling rate" is the number of empirical samples used in negative edge sampling (samples from $p_{\mathrm{ns}}$). For each hyperparameter, the center value is the default. To control computational cost, one hyperparameter was varied at a time, with all others set to their default values. The performance of GLUE was robust across a wide range of hyperparameter settings, except for failed alignments in which the adversarial alignment weight was too low or no hidden layers were used in the neural networks (equivalently a linear model with insufficient capacity).

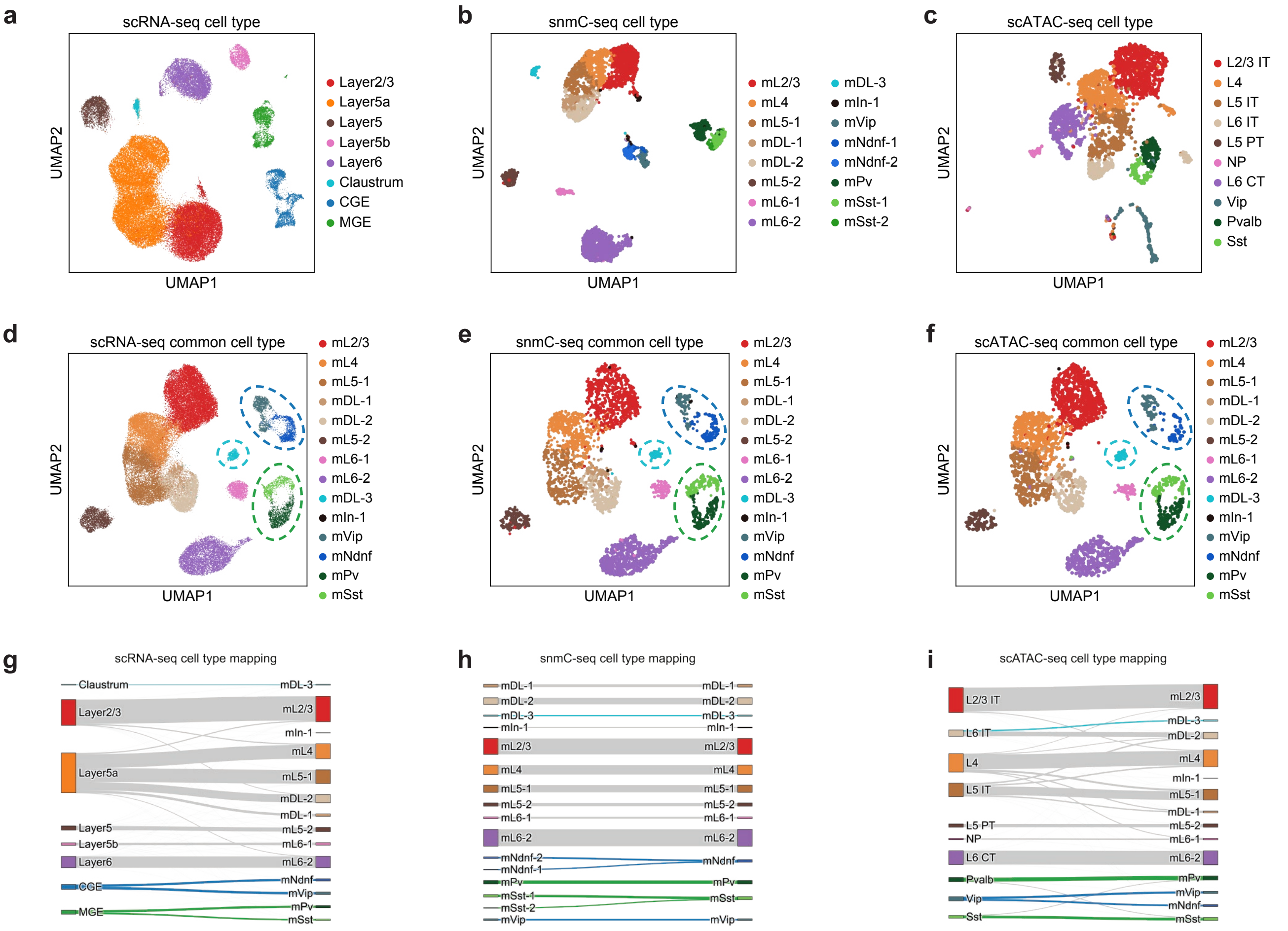


**Supplementary Fig. 5 Triple-omics alignment and label transfer.**
**a**-**c**, UMAP visualizations of the cell embeddings before integration for **a**, scRNA-seq, **b**, snmC-seq, and **c**, scATAC-seq, colored by the original cell types. **d**-**f**, UMAP visualizations of the integrated cell embeddings for **d**, scRNA-seq, **e**, snmC-seq, and **f**, scATAC-seq, colored by the unified cell types (labels transferred from snmC-seq). **g**-**i**, Alluvial diagrams comparing the original cell types and unified cell types for **g**, scRNA-seq, **h**, snmC-seq, and **i**, scATAC-seq. The original cell types are to the left, and the unified cell types are to the right. Cells relabeled to "mPv" and "mSst" are highlighted with green flows. Cells relabeled to "mNdnf" and "mVip" are highlighted with dark blue flows. Cells relabeled to "mDL-3" are highlighted with light blue flows.

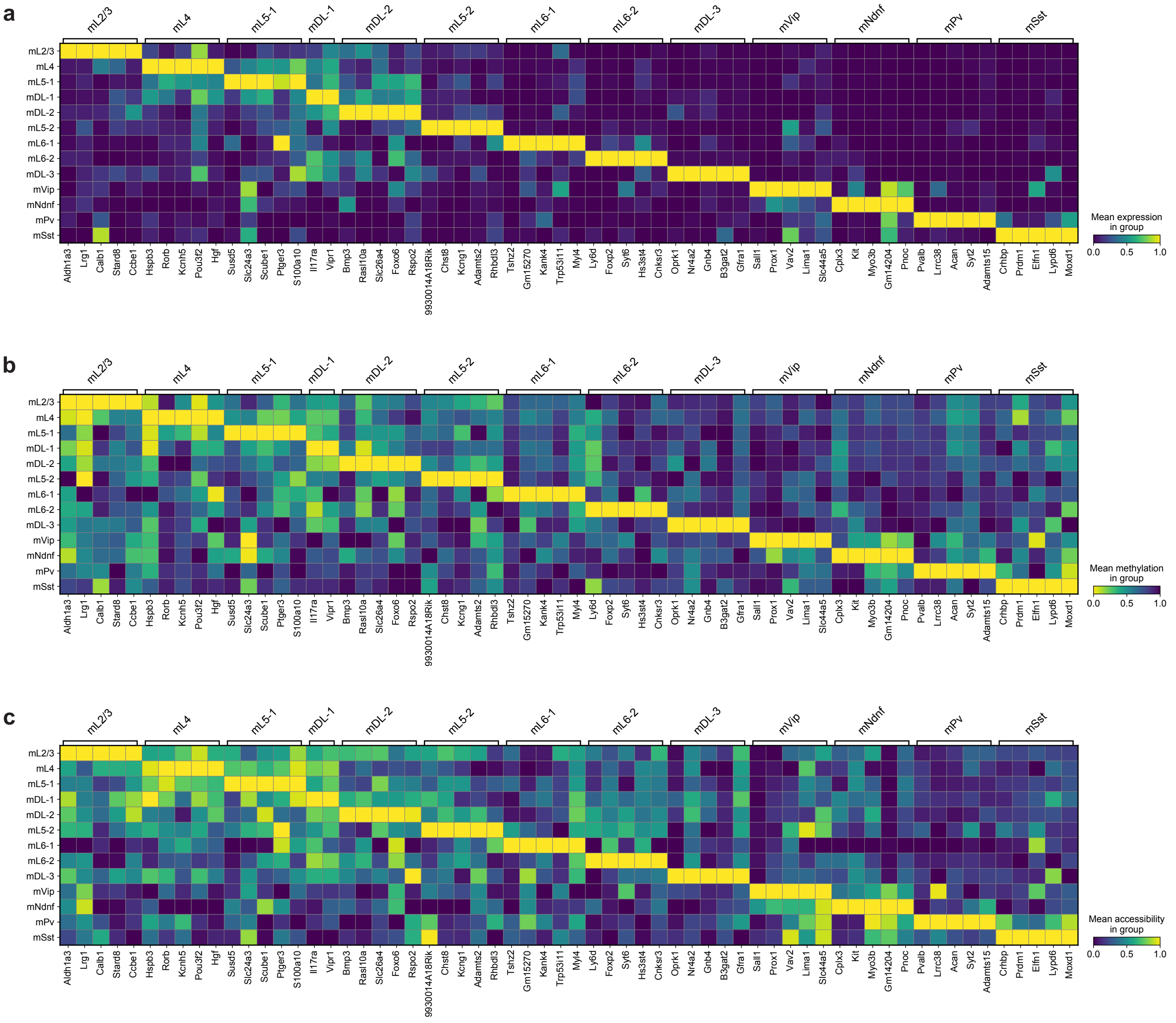


**Supplementary Fig. 6 Consensus cell type markers across three omics layers.**
**a**, Expression in scRNA-seq. **b**, Gene body DNA methylation in snmC-seq. **c**, Chromatin accessibility in scATAC-seq. Note that an inverted color bar is used for DNA methylation.

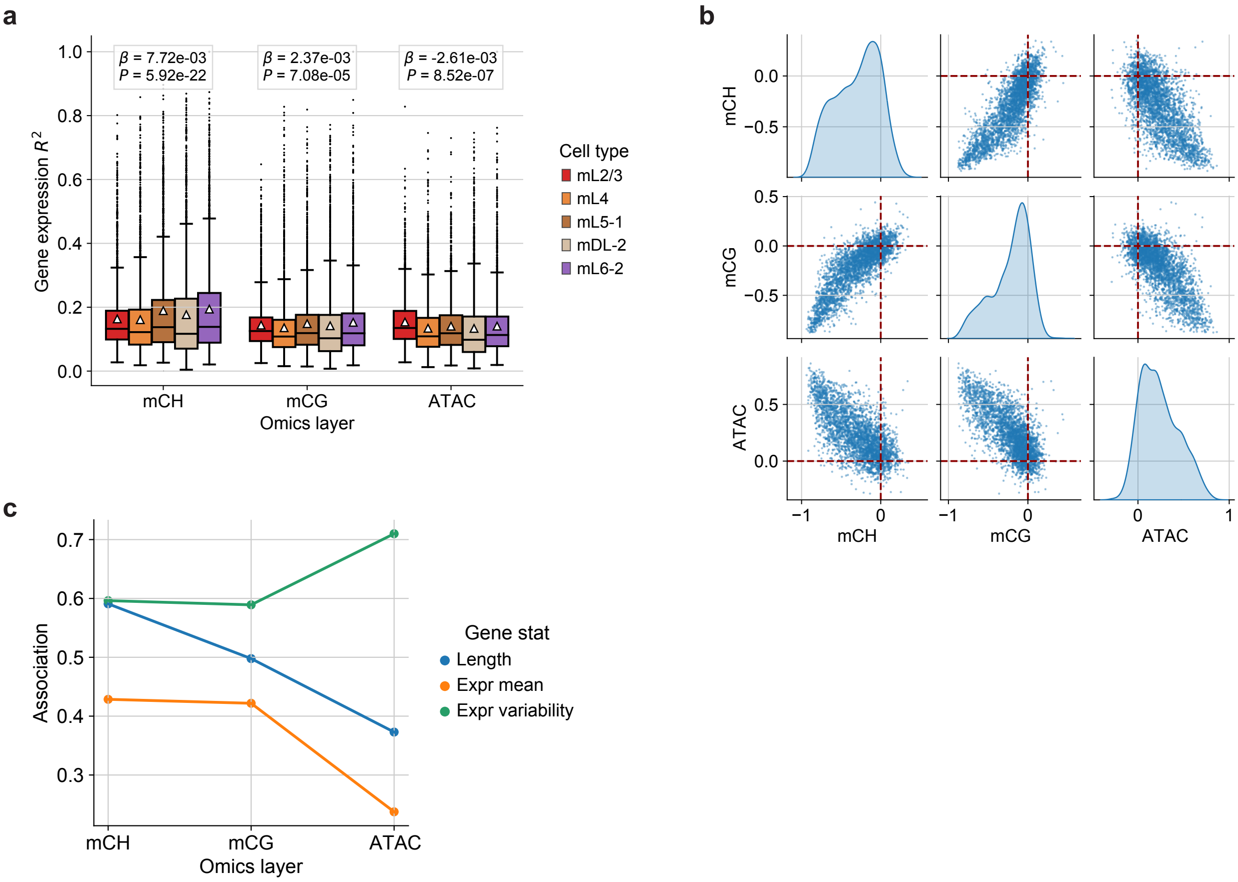


**Supplementary Fig. 7 Epigenetic contributions to gene expression.**
**a**, Coefficient of determination ($R^2$) for predicting gene expression based on each epigenetic layer in different cell types. The box plots indicate the medians (centerlines), means (triangles), 1st and 3rd quartiles (hinges), and minima and maxima (whiskers). Above each epigenetic layer, the linear regression slope ($\beta$) and its $P$ value are displayed, with the cell type as the regressor. **b**, Pearson's correlations between different epigenetic states and gene expression. **c**, Associations (defined as the absolute values of Pearson's correlations) between the correlations in **b** and different gene characteristics.

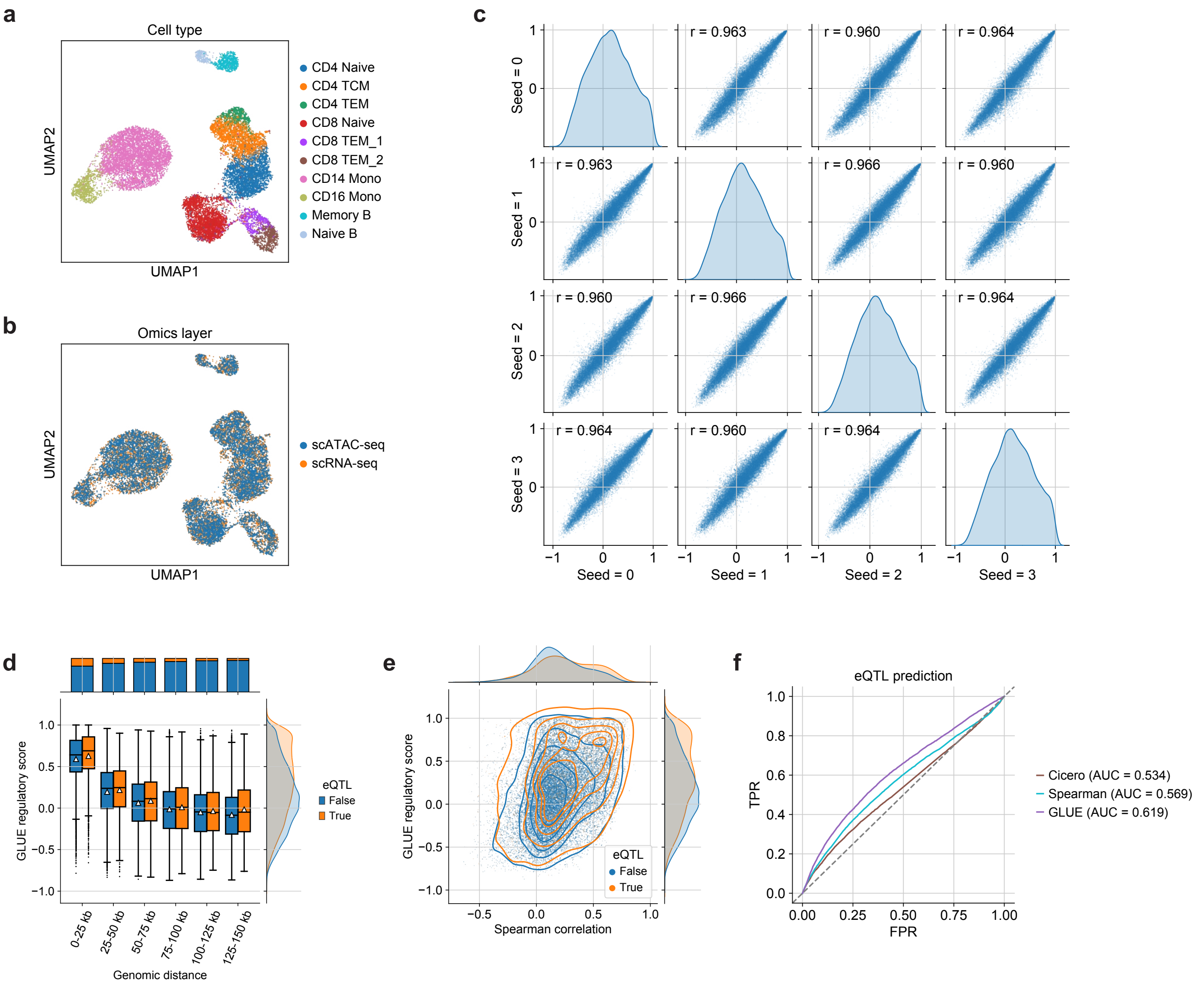


### Supplementary Fig. 8 Model-based regulatory inference using distance-based power-law interactions as guidance.

**a**, **b**, UMAP visualizations of the integrated cell embeddings colored by **a**, cell types, and **b**, omics layers. **c**, Pearson's correlation coefficients for the GLUE regulatory scores across different random seeds. **d**, GLUE regulatory scores for peak-gene pairs across different genomic ranges, grouped by whether they had eQTL support. The box plots indicate the medians (centerlines), means (triangles), 1st and 3rd quartiles (hinges), and minima and maxima (whiskers). **e**, Comparison between the GLUE regulatory scores and the empirical peak-gene correlations computed on paired cells. Peak-gene pairs are colored by whether they had eQTL support. **f**, ROC (receiver operating characteristic) curves for predicting eQTL interactions based on different peak-gene association scores.

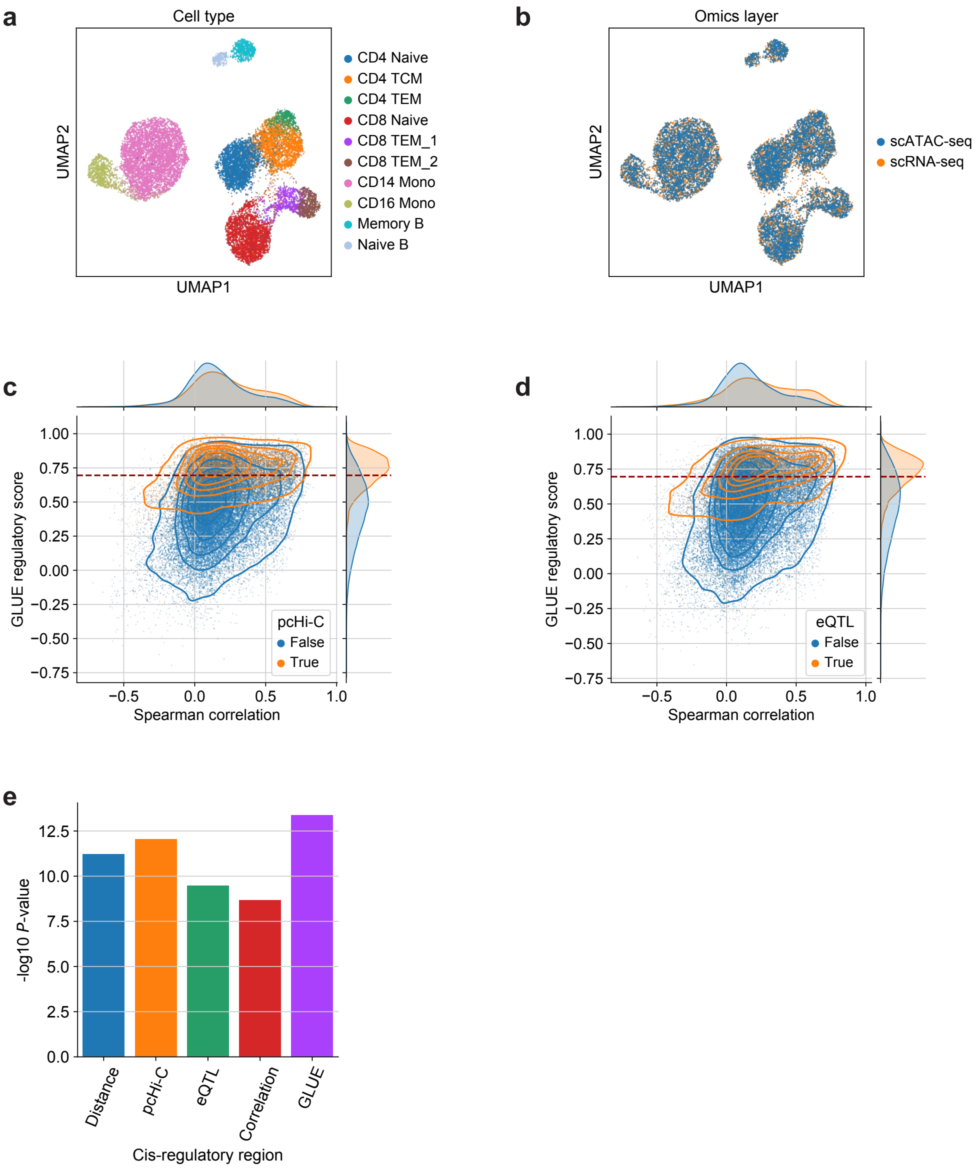


**Supplementary Fig. 9 Model-based regulatory inference using a combination of distance-based, eQTL and pcHi-C interactions as guidance.**

**a**, **b**, UMAP visualizations of the integrated cell embeddings colored by **a**, cell types, and **b**, omics layers. **c**, **d**, Comparison between the GLUE regulatory scores and the empirical peak-gene correlations computed on paired cells. The peak-gene pairs are colored by whether they had **c**, pcHi-C support or **d**, eQTL support. Red dashed lines indicate the 75th percentile of the GLUE regulatory scores, which was used as a cutoff. The GLUE-identified interactions mostly exhibited positive empirical correlations and covered the majority of pcHi-C and eQTL support, while the selection of peak-gene pairs based solely on empirical correlation would lead to much lower external support. **e**, Consistency between the TF-target gene networks constructed with different peak-gene association methods and the manually curated connections in the TRRUST v2 database (Fisher's exact test).

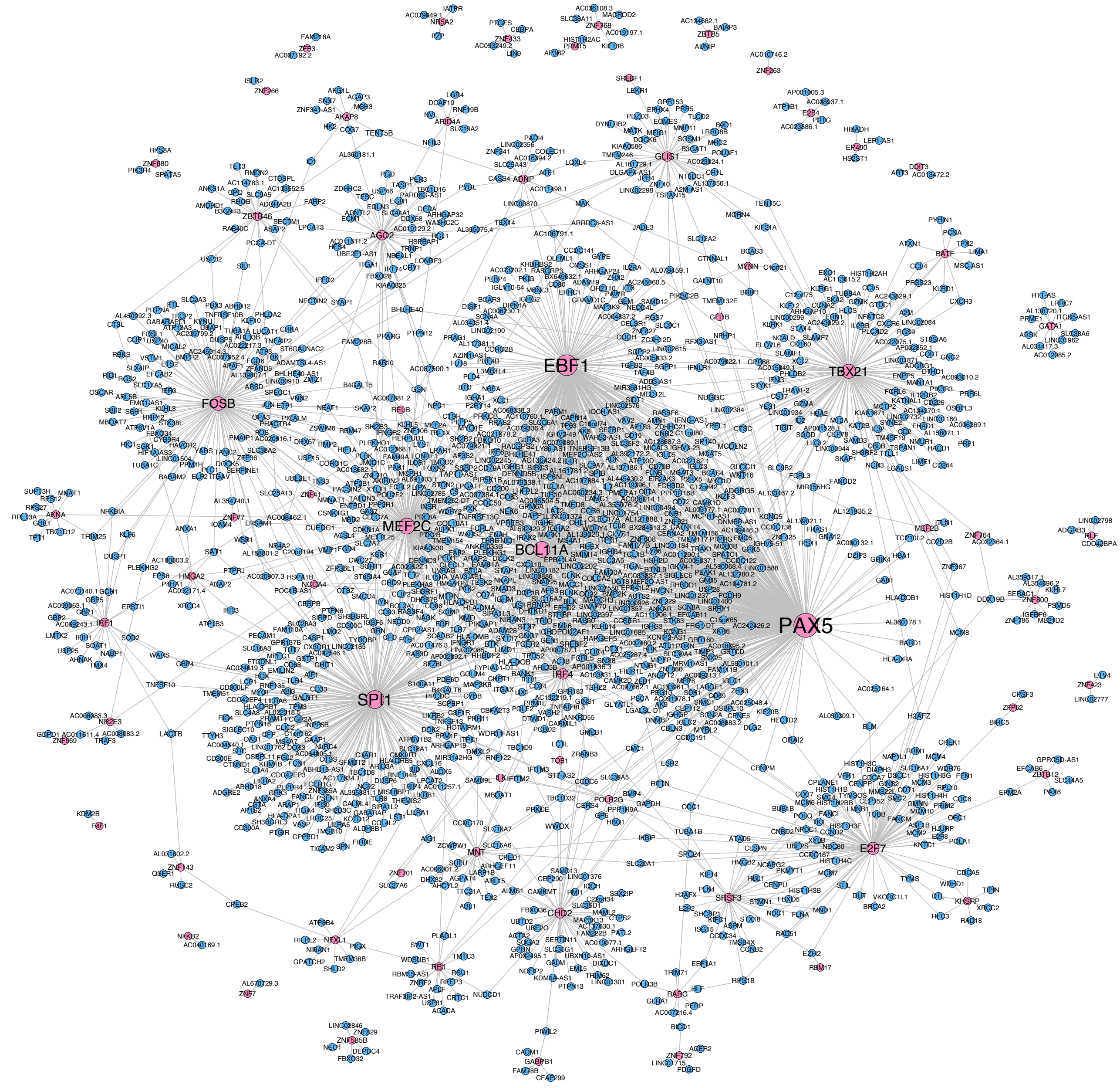


**Supplementary Fig. 10 Inferred TF-target gene regulatory network in PBMC.**

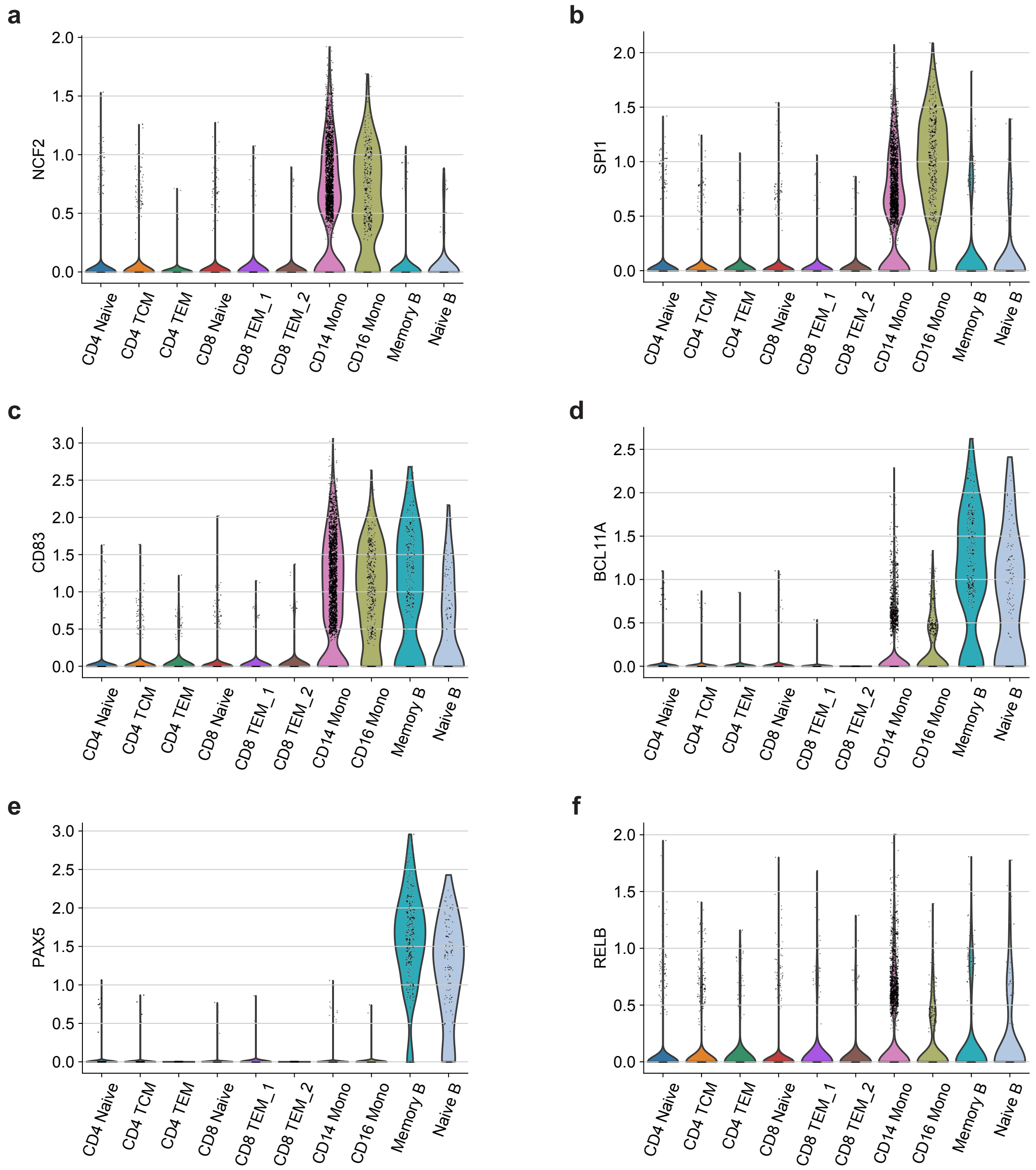


**Supplementary Fig. 11 TF-target gene expression levels in different cell types.**
**a**, **b**, Expression levels of **a**, *NCF2*, and its regulator **b**, *SPI1* in different cell types. **c-f**, Expression levels of **c**, *CD83*, and its inferred regulators **d**, *BCL11A*, **e**, *PAX5*, and **f**, *RELB* in different cell types.

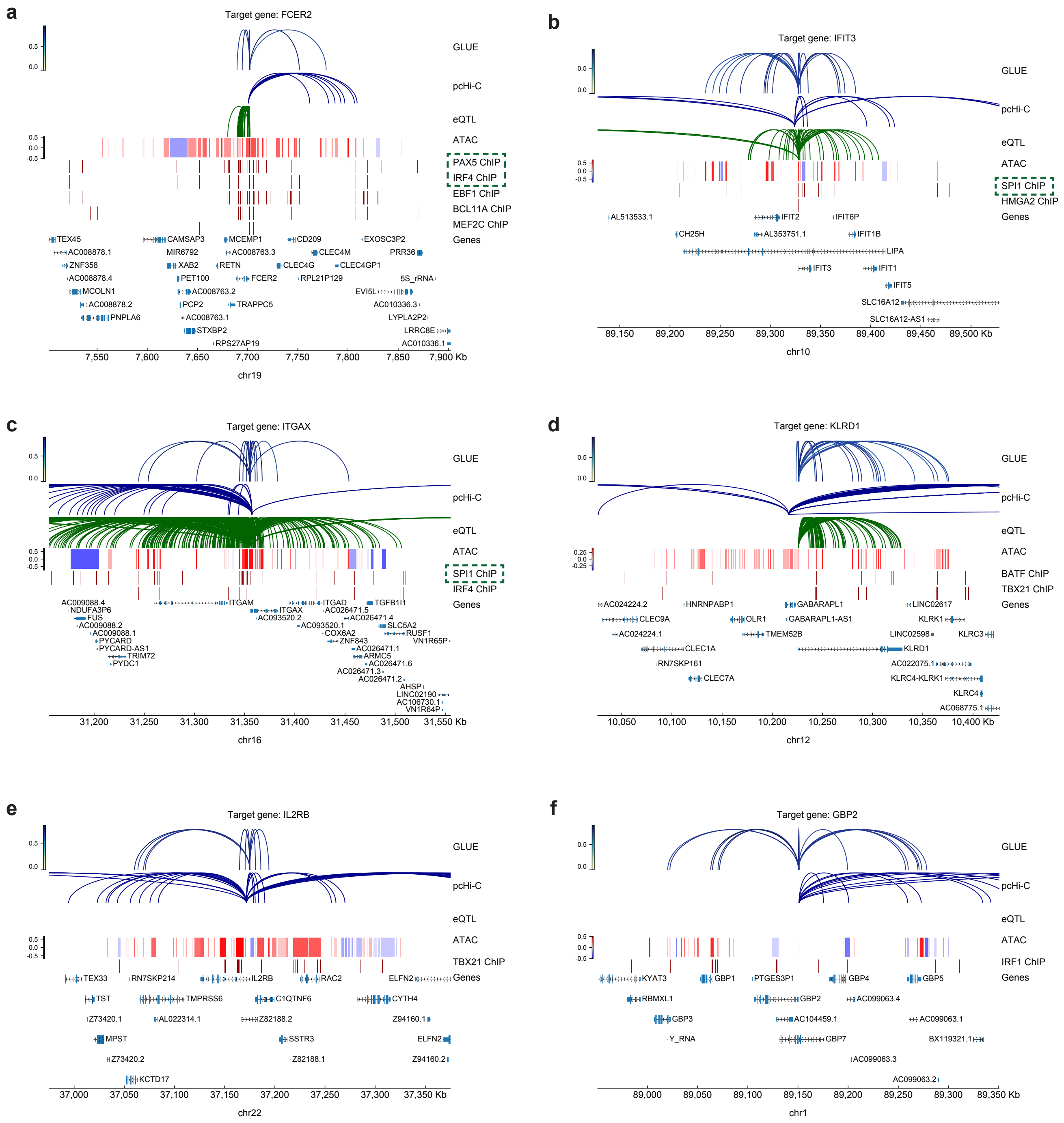


**Supplementary Fig. 12 Regulatory inference and evidence per gene.**
GLUE-identified cis-regulatory interactions for **a**, *FCER2*, **b**, *IFIT3*, **c**, *ITGAX*, **d**, *KLRD1*, **e**, *IL2RB*, and **f**, *GBP2*, along with individual pieces of regulatory evidence. For each gene, the ChIP-seq tracks correspond to inferred TF regulators. Known regulators are highlighted with green boxes.

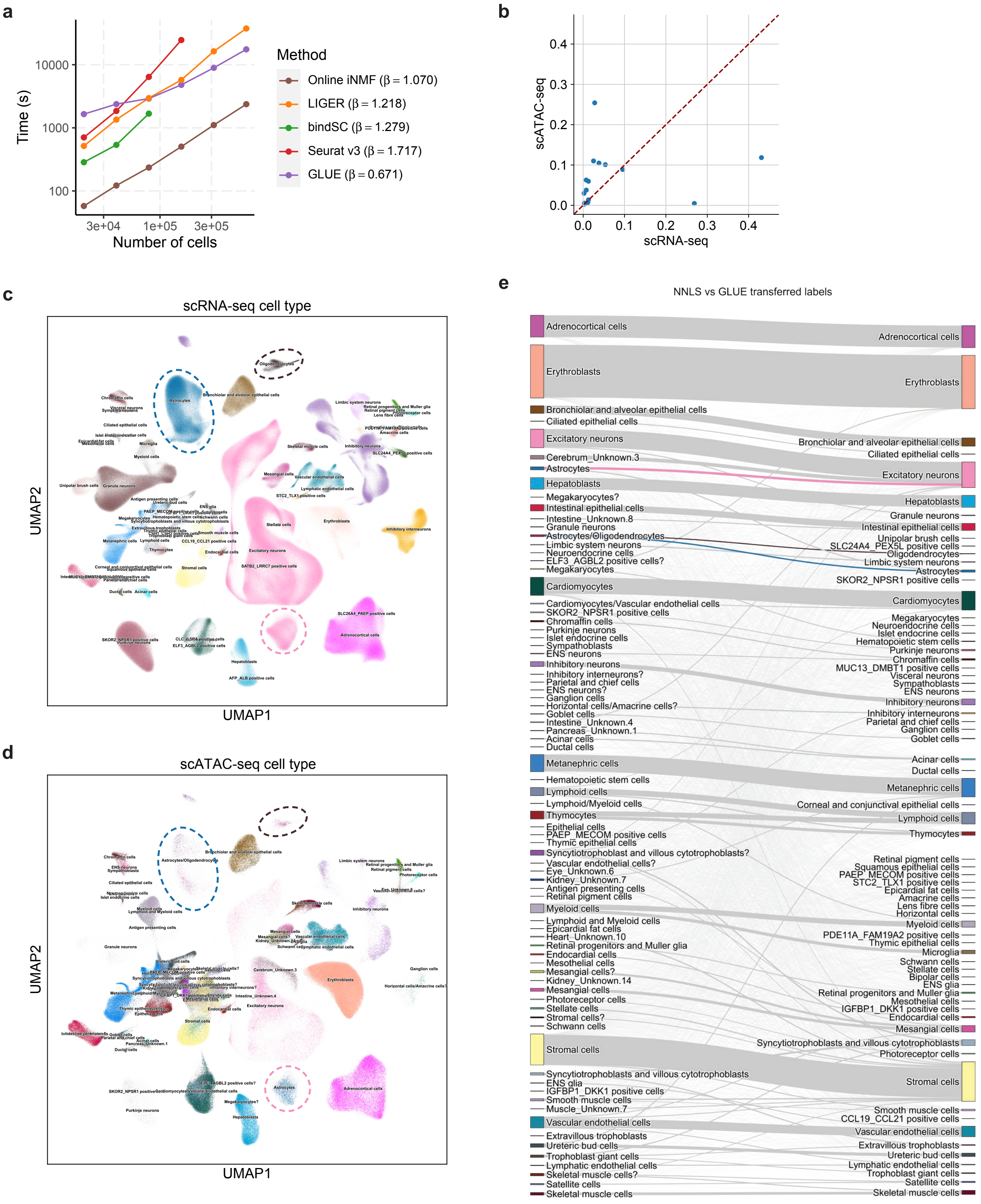
a
Time (s)
10000
1000
100
3e+04
1e+05
3e+05
Number of cells
Method
Online iNMF (β = 1.070)
LIGER (β = 1.218)
bindSC (β = 1.279)
Seurat v3 (β = 1.717)
GLUE (β = 0.671)
b
scATAC-seq
scRNA-seq
0.0
0.1
0.2
0.3
0.4
c
scRNA-seq cell type
UMAP1
UMAP2
d
scATAC-seq cell type
UMAP1
UMAP2
e
NNLS vs GLUE transferred labels
Adrenocortical cells
Erythroblasts
Bronchiolar and alveolar epithelial cells
Ciliated epithelial cells
Excitatory neurons
Cerebrum_Unknown.3
Astrocytes
Hepatoblasts
Megakaryocytes?
Intestinal epithelial cells
Intestine_Unknown.8
Granule neurons
Astrocytes/Oligodendrocytes
Limbic system neurons
Neuroendocrine cells
ELF3_AGBL2 positive cells?
Megakaryocytes
Cardiomyocytes
Cardiomyocytes/Vascular endothelial cells
SKOR2_NPSR1 positive cells
Chromaffin cells
Purkinje neurons
Islet endocrine cells
Sympathoblasts
ENS neurons
Inhibitory neurons
Inhibitory interneurons?
Parietal and chief cells
ENS neurons?
Ganglion cells
Horizontal cells/Amacrine cells?
Goblet cells
Intestine_Unknown.4
Pancreas_Unknown.1
Acinar cells
Ductal cells
Metanephric cells
Hematopoietic stem cells
Lymphoid cells
Lymphoid/Myeloid cells
Thymocytes
Epithelial cells
PAEP_MECOM positive cells
Thymic epithelial cells
Syncytiotrophoblasts and villous cytotrophoblasts?
Vascular endothelial cells?
Eye_Unknown.6
Kidney_Unknown.7
Antigen presenting cells
Retinal pigment cells
Myeloid cells
Lymphoid and Myeloid cells
Epicardial fat cells
Heart_Unknown.10
Retinal progenitors and Muller glia
Endocardial cells
Mesothelial cells
Mesangial cells?
Kidney_Unknown.14
Mesangial cells
Photoreceptor cells
Stellate cells
Stromal cells?
Schwann cells
Stromal cells
Syncytiotrophoblasts and villous cytotrophoblasts
ENS glia
IGFBP1_DKK1 positive cells
Smooth muscle cells
Muscle_Unknown.7
Vascular endothelial cells
Extravillous trophoblasts
Ureteric bud cells
Trophoblast giant cells
Lymphatic endothelial cells
Skeletal muscle cells?
Satellite cells
Skeletal muscle cells
Adrenocortical cells
Erythroblasts
Bronchiolar and alveolar epithelial cells
Ciliated epithelial cells
Excitatory neurons
Hepatoblasts
Granule neurons
Intestinal epithelial cells
Unipolar brush cells
SLC24A4_PEX5L positive cells
Oligodendrocytes
Limbic system neurons
Astrocytes
SKOR2_NPSR1 positive cells
Cardiomyocytes
Megakaryocytes
Neuroendocrine cells
Islet endocrine cells
Hematopoietic stem cells
Purkinje neurons
Chromaffin cells
MUC13_DMBT1 positive cells
Visceral neurons
Sympathoblasts
ENS neurons
Inhibitory neurons
Inhibitory interneurons
Parietal and chief cells
Ganglion cells
Goblet cells
Acinar cells
Ductal cells
Metanephric cells
Corneal and conjunctival epithelial cells
Lymphoid cells
Thymocytes
Retinal pigment cells
Squamous epithelial cells
PAEP_MECOM positive cells
STC2_TLX1 positive cells
Epicardial fat cells
Amacrine cells
Lens fibre cells
Horizontal cells
Myeloid cells
PDE11A_FAM19A2 positive cells
Thymic epithelial cells
Microglia
Schwann cells
Stellate cells
Bipolar cells
ENS glia
Retinal progenitors and Muller glia
Mesothelial cells
IGFBP1_DKK1 positive cells
Endocardial cells
Mesangial cells
Syncytiotrophoblasts and villous cytotrophoblasts
Photoreceptor cells
Stromal cells
Smooth muscle cells
CCL19_CCL21 positive cells
Vascular endothelial cells
Extravillous trophoblasts
Ureteric bud cells
Lymphatic endothelial cells
Trophoblast giant cells
Satellite cells
Skeletal muscle cells

**Supplementary Fig. 13 Scalability benchmarking and integration of a multi-omics human cell atlas.**

**a**, Time costs of different methods on subsampled atlases of varying sizes. The time costs and cell numbers are both plotted in log-scale, so the slope $\beta$ represents the exponent in original scale, i.e., $\beta = 2$ represents quadratic scalability, $\beta = 1$ represents linear scalability, and $\beta < 1$ represents sublinear scalability. While online iNMF was the fastest method on the tested data sizes, GLUE has a lower $\beta$, and could surpass online iNMF as data size increases to dozens of millions. **b**, Organ compositions in scRNA-seq and scATAC-seq. **c**, UMAP visualizations of the integrated cell embeddings showing only the scRNA-seq cells, colored by the original cell types. **d**, UMAP visualizations of the integrated cell embeddings showing only the scATAC-seq cells, colored by the original cell types. **e**, Alluvial diagram comparing the NNLS (non-negative least squares)-based and GLUE-based cell type annotations. The original NNLS-based annotations are to the left, and the GLUE-based annotations are to the right. Cells originally labeled as "Astrocytes" but mapped to "Excitatory neurons" are highlighted with pink flows. Cells originally labeled as "Astrocytes/Oligodendrocytes" but mapped to "Astrocytes" are highlighted with blue flows. Cells originally labeled as "Astrocytes/Oligodendrocytes" but mapped to "Oligodendrocytes" are highlighted with brown flows.

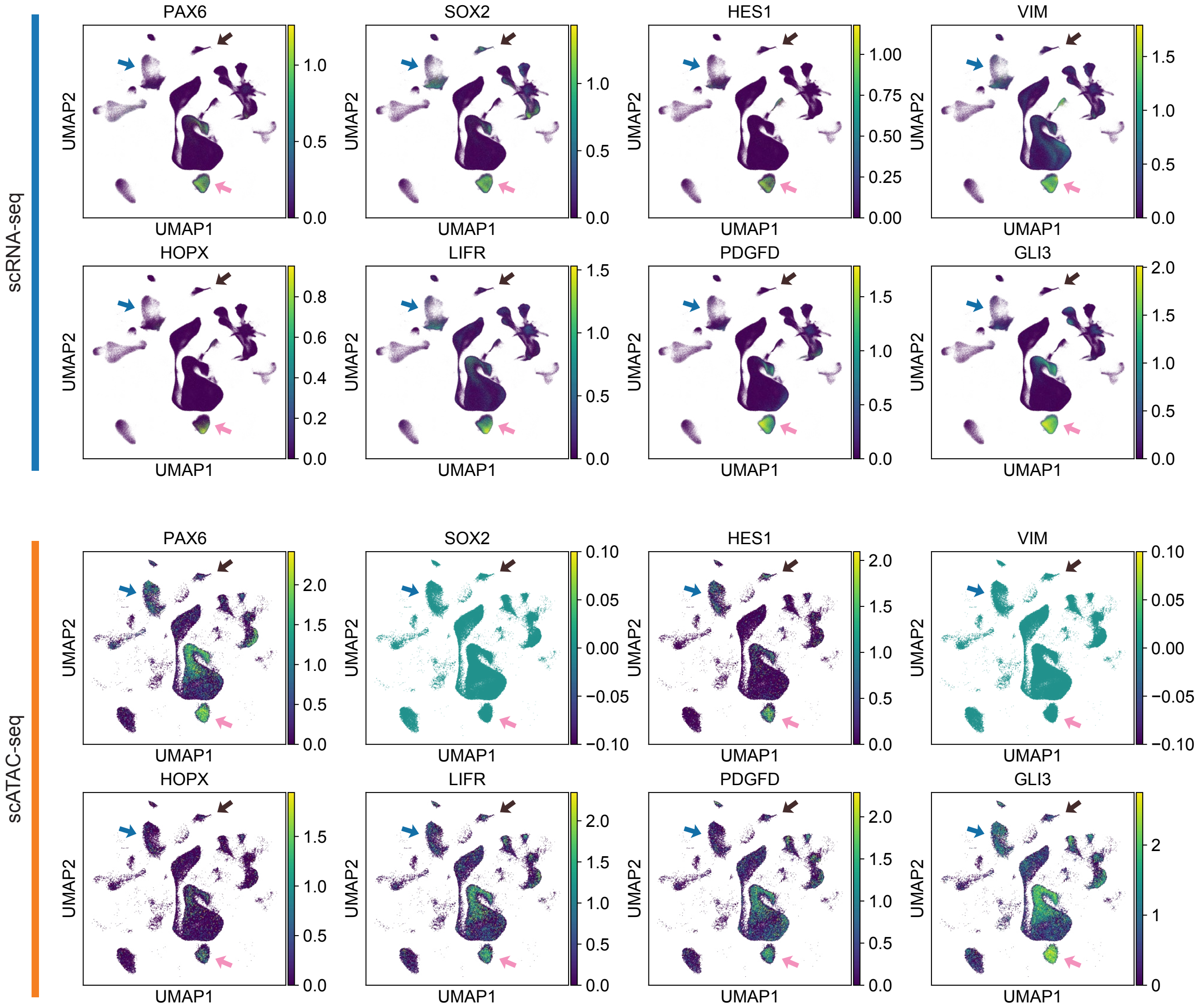


**Supplementary Fig. 14 Gene expression and chromatin accessibility patterns of neural progenitor markers in cerebrum cells.**

Putative neural progenitors are highlighted with pink arrows, astrocytes are highlighted with blue arrows, and oligodendrocytes are highlighted with brown arrows. *SOX2* and *VIM* were not detected in scATAC-seq, due to limitations in ATAC gene-level score calculation (no peaks overlap gene body and 2 kb upstream from TSS).

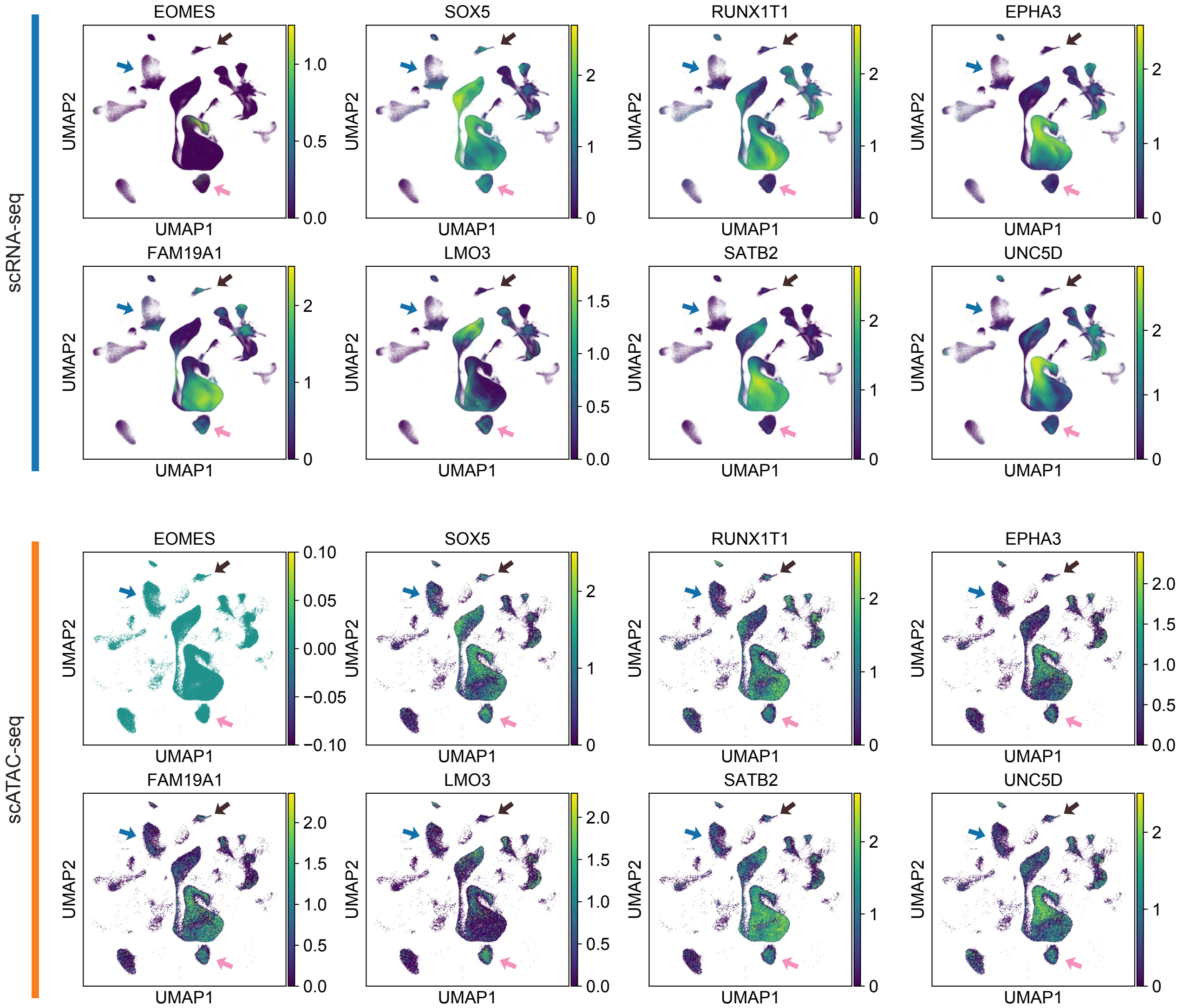


**Supplementary Fig. 15 Gene expression and chromatin accessibility patterns of excitatory neuron markers in cerebrum cells.**

Putative neural progenitors are highlighted with pink arrows, astrocytes are highlighted with blue arrows, and oligodendrocytes are highlighted with brown arrows. *EOMES* was not detected in scATAC-seq, due to limitations in ATAC gene-level score calculation (no peaks overlap gene body and 2 kb upstream from TSS).

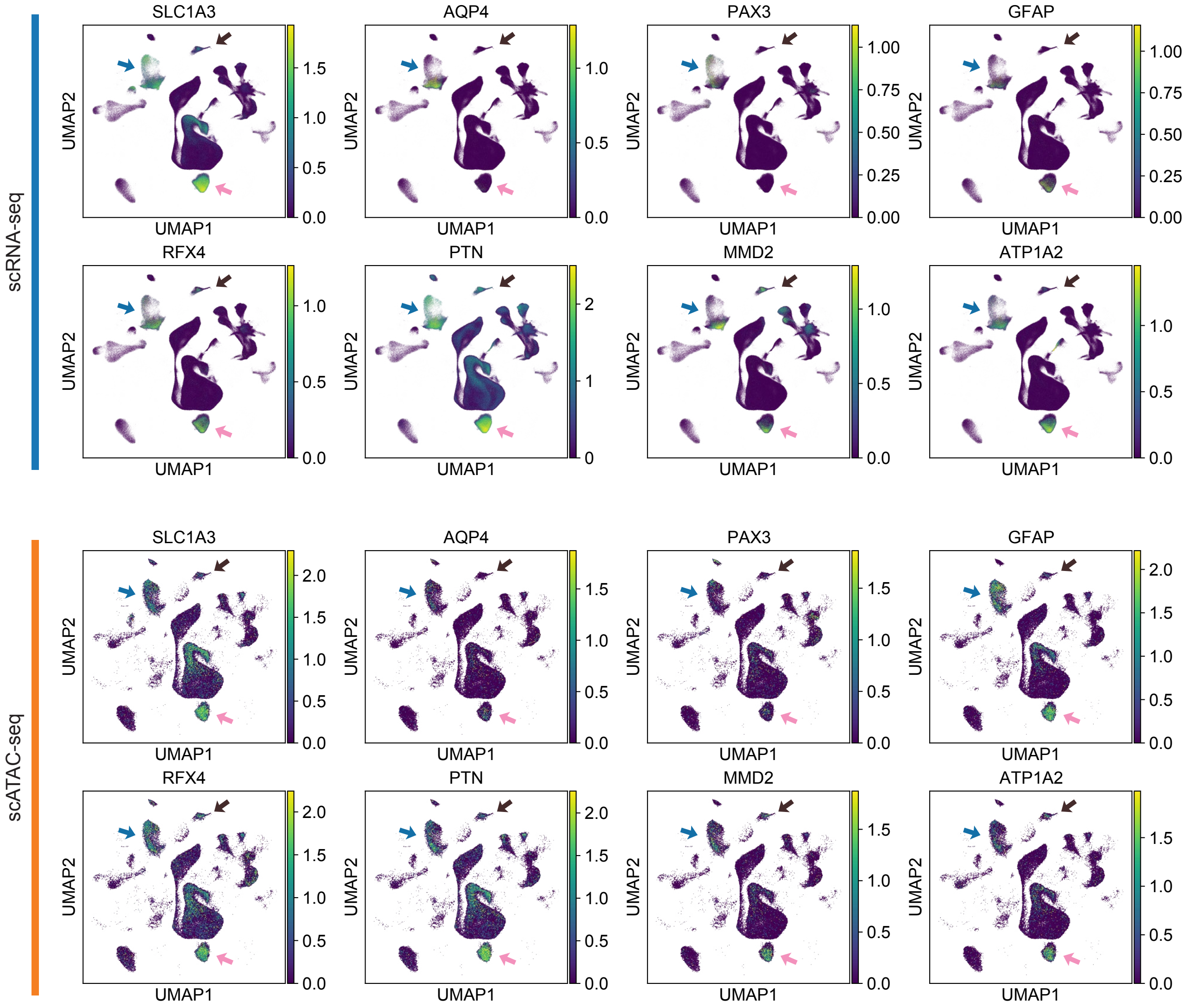


**Supplementary Fig. 16 Gene expression and chromatin accessibility patterns of astrocyte markers in cerebrum cells.**

Putative neural progenitors are highlighted with pink arrows, astrocytes are highlighted with blue arrows, and oligodendrocytes are highlighted with brown arrows.

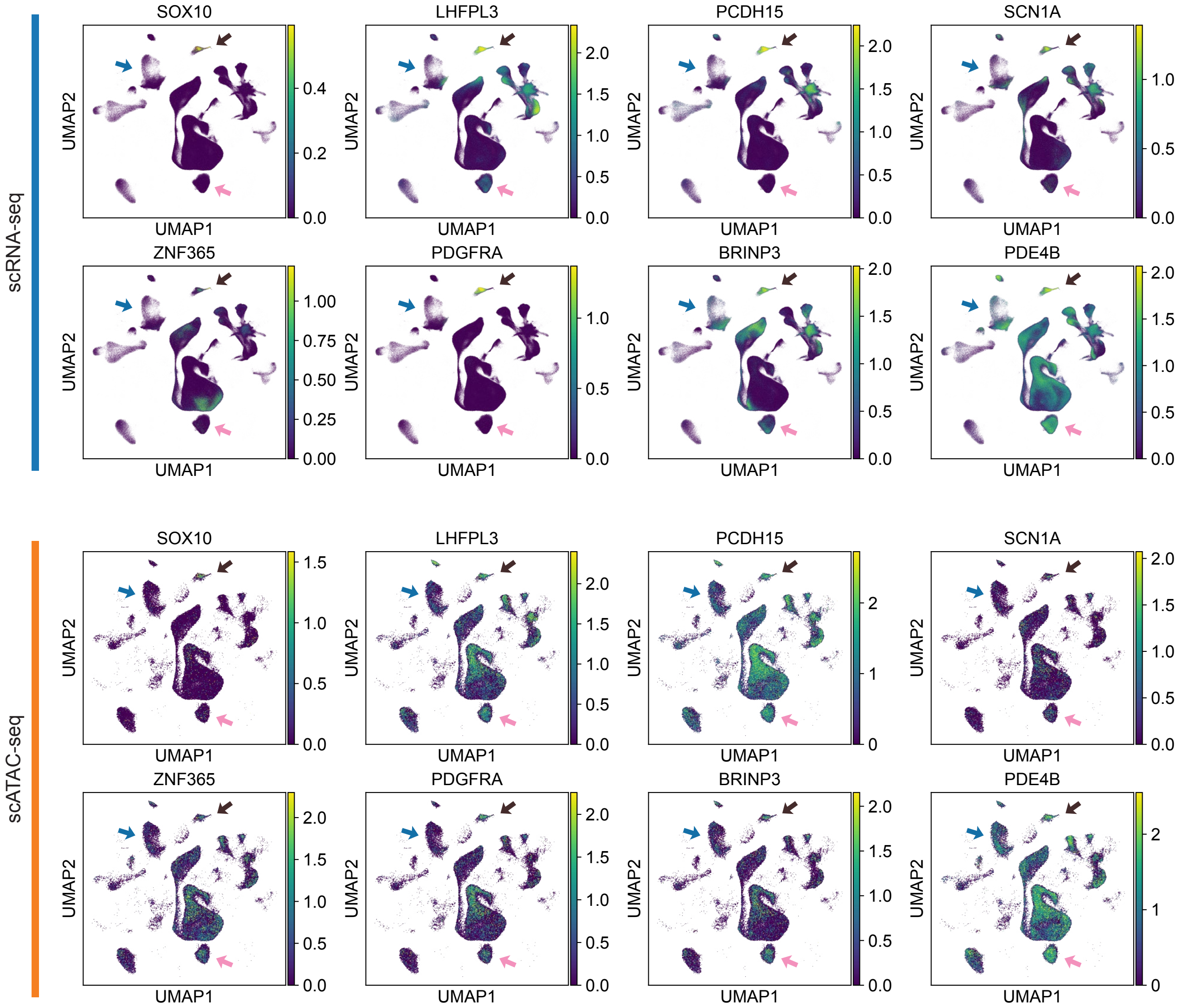


**Supplementary Fig. 17 Gene expression and chromatin accessibility patterns of oligodendrocyte markers in cerebrum cells.**
Putative neural progenitors are highlighted with pink arrows, astrocytes are highlighted with blue arrows, and oligodendrocytes are highlighted with brown arrows.

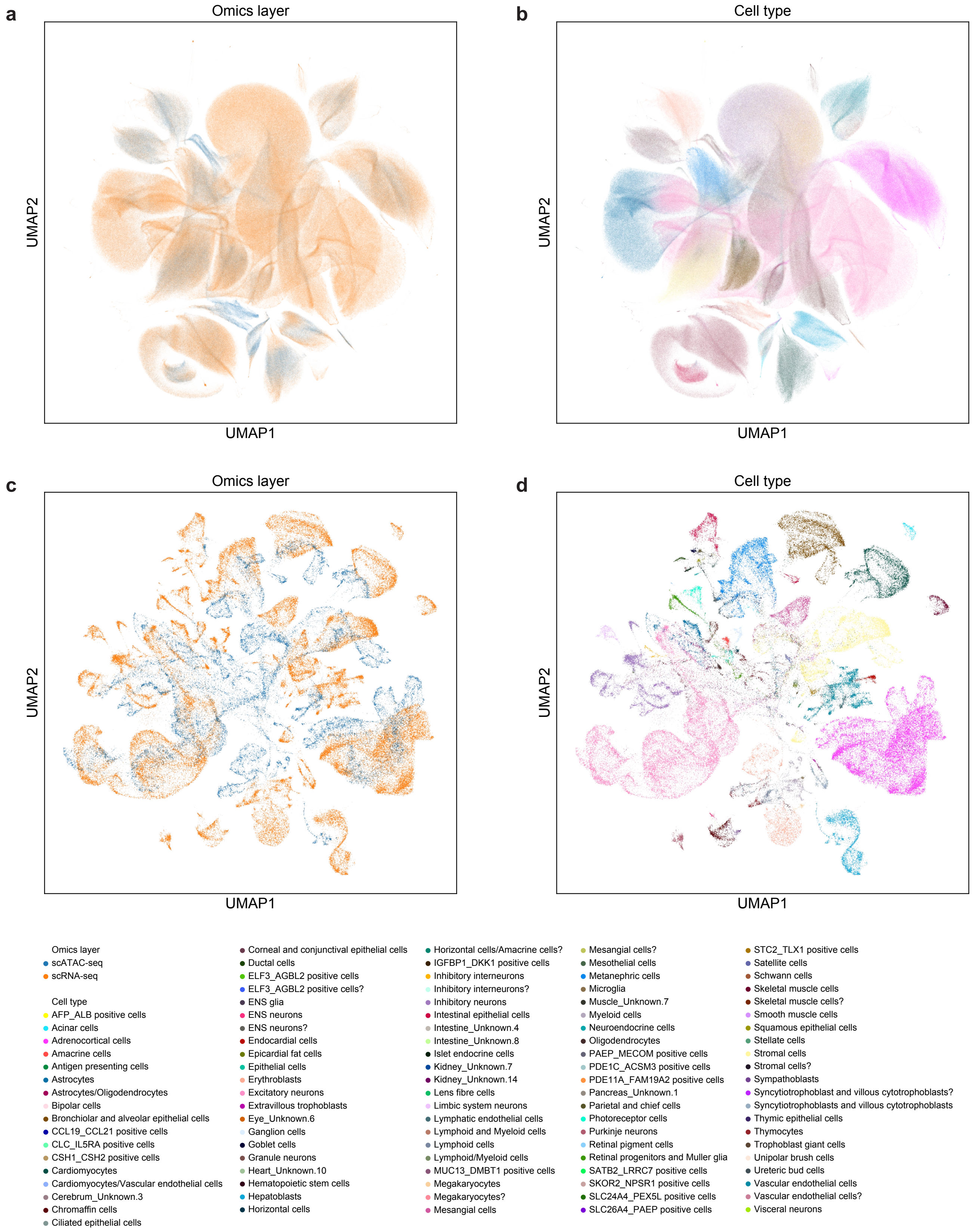


**Supplementary Fig. 18 UMAP visualization of the multi-omics human cell atlas integrated by other methods.**
**a**, **b**, Online iNMF-integrated cell embeddings colored by **a**, omics layers, and **b**, cell types. **c**, **d**, Seurat v3-integrated cell embeddings of aggregated metacells colored by **c**, omics layers, and **d**, cell types.

**Supplementary Table 1 Public datasets used in the study.**

**Supplementary Table 2 Detailed benchmark data.**

**Supplementary Table 3 Regulatory interactions in the GLUE-derived TF-target gene network.**
